# Metabolic pathway prediction using non-negative matrix factorization with improved precision

**DOI:** 10.1101/2020.05.27.119826

**Authors:** Abdur Rahman M. A. Basher, Ryan J. McLaughlin, Steven J. Hallam

## Abstract

Machine learning provides a probabilistic framework for metabolic pathway inference from genomic sequence information at different levels of complexity and completion. However, several challenges including pathway features engineering, multiple mapping of enzymatic reactions and emergent or distributed metabolism within populations or communities of cells can limit prediction performance. In this paper, we present triUMPF, <u>tri</u>ple non-negative matrix factorization (NMF) with comm<u>u</u>nity detection for <u>m</u>etabolic <u>p</u>athway in<u>f</u>erence, that combines three stages of NMF to capture myriad relationships between enzymes and pathways within a graph network. This is followed by community detection to extract higher order structure based on the clustering of vertices which share similar statistical properties. We evaluated triUMPF performance using experimental datasets manifesting diverse multi-label properties, including Tier 1 genomes from the BioCyc collection of organismal Pathway/Genome Databases and low complexity microbial communities. Resulting performance metrics equaled or exceeded other prediction methods on organismal genomes with improved precision on multi-organismal datasets.

**Availability and implementation:** The software package, and installation instructions are published on github.com/triUMPF

## 1 Introduction

Pathway reconstruction from genomic sequence information is an essential step in describing the metabolic potential of cells at the individual, population and community levels of biological organization [12, 18, 25]. Resulting pathway representations provide a foundation for defining regulatory processes, modeling metabolite flux and engineering cells and cellular consortia for defined process outcomes [11, 20]. The integral nature of the pathway prediction problem has prompted both gene-centric e.g. mapping annotated proteins onto known pathways using a reference database based on sequence homology, and heuristic or rule-based pathway-centric approaches including PathoLogic [16] and MinPath [38]. In parallel, the development of trusted sources of curated metabolic pathway information including the Kyoto Encyclopedia of Genes and Genomes (KEGG) [15] and MetaCyc [4] provides training data for the design of more flexible machine learning (ML) algorithms for pathway inference. While ML approaches have been adopted widely in metabolomics research [3,34] they have gained less traction when applied to predicting pathways directly from annotated gene lists.

Dale and colleagues conducted the first in-depth exploration of ML approaches for pathway prediction using Tier 1 (T1) organismal Pathway/Genome Databases (PGDB) [5] from the BioCyc collection randomly divided into training and test sets [7]. Features were developed based on rule-sets used by the PathoLogic algorithm in Pathway Tools to construct PGDBs [16]. Resulting performance metrics indicated that standard ML approaches rivaled PathoLogic performance with the added benefit of probability scores [7]. More recently Basher and colleagues developed <u>*m*</u>ulti-<u>*l*</u>abel based on <u>*l*</u>ogistic re<u>*g*</u>ression for <u>*p*</u>athway p<u>*r*</u>ediction (mlLGPR), a multi-label classification approach that uses logistic regression and feature vectors inspired by the work of Dale and colleagues to predict metabolic pathways from genomic sequence information at different levels of complexity and completion [25].

Although mlLGPR performed effectively on organismal genomes, pathway prediction outcomes for multi-organismal datasets were less optimal due in part to missing or noisy feature information. In an effort to solve this problem, Basher and Hallam evaluated the use of representational learning methods to learn a neural embedding-based low-dimensional space of metabolic features based on a three-layered network architecture consisting of compounds, enzymes, and pathways [24]. Learned feature vectors improved pathway prediction performance on organismal genomes and motivated the use of graphical models for multi-organismal features engineering.

Here we describe <u>tri</u>ple non-negative matrix factorization (NMF) with comm<u>u</u>nity detection for <u>m</u>etabolic <u>p</u>athway in<u>f</u>erence (triUMPF) combining three stages of NMF to capture relationships between enzymes and pathways within a network [9] followed by community detection to extract higher order network structure [8]. Non-negative matrix factorization is a data reduction and exploration method in which the original and factorized matrices have the property of non-negative elements with reduced ranks or features [9]. In contrast to other dimension reduction methods, such as principal component analysis [2], NMF both reduces the number of features and preserves information needed to reconstruct the original data [37]. This has important implications for noise robust feature extraction from sparse matrices including datasets associated with gene expression analysis and pathway prediction [37].

For pathway prediction, triUMPF uses three graphs, one representing associations between pathways and enzymes indicated by enzyme commission (EC)) numbers [1], one representing interactions between enzymes and another representing interactions between pathways. The two interaction graphs adopt the *subnetworks* concept introduced in BiomeNet [32] and MetaNetSim [14], where a subnetwork is a linked series of connected nodes (e.g. reactions and pathways). In the literature, a subnetwork is commonly referred to as a *community* [30], which defines a set of densely connected nodes within a subnetwork. It is important to emphasize that unless otherwise indicated, the use of the term community in this work refers to a subnetwork community based on statistical properties of a network rather than a community of organisms. Community detection is performed on both interaction graphs (pathways and enzymes) to identify subnetworks among pathways.

We evaluated triUMPF’s prediction performance in relation to other methods including MinPath, PathoLogic, and mlLGPR on a set of T1 PGDBs, low complexity microbial communities including symbiont genomes encoding distributed metabolic pathways for amino acid biosynthesis [26], genomes used in the Critical Assessment of Metagenome Interpretation (CAMI) initiative [31], and whole genome shotgun sequences from the Hawaii Ocean Time Series (HOTS) [33] following information hierarchy-based benchmarks initially developed for mlLGPR enabling more robust comparison between pathway prediction methods [25].

## 2 Methods

In this section, we provide a general description of triUMPF components, presented in Fig. 1. At the very beginning, MetaCyc is applied to: i)- extract three association matrices, indicated in step Fig. 1(a), one representing associations between pathways and enzymes (P2E) indicated by enzyme commission (EC)) numbers [27], one representing interactions between enzymes (E2E) and another representing interactions between pathways (P2P), and ii)- automatically generate features corresponding pathways and enzymes (or EC) from pathway2vec [24] in Fig. 1(b). Then, triUMPF is trained in two phases: i)- decomposition of the pathway EC association matrix in Fig. 1(c), and ii)- subnetwork or community reconstruction while, simultaneously, learning optimal multi-label pathway parameters in Figs 1(d-f). Below, we discuss these two phases while the analytical expressions of triUMPF are explained in Appx. Sections 5.1, 5.2, and 5.3.

**Figure 1:**
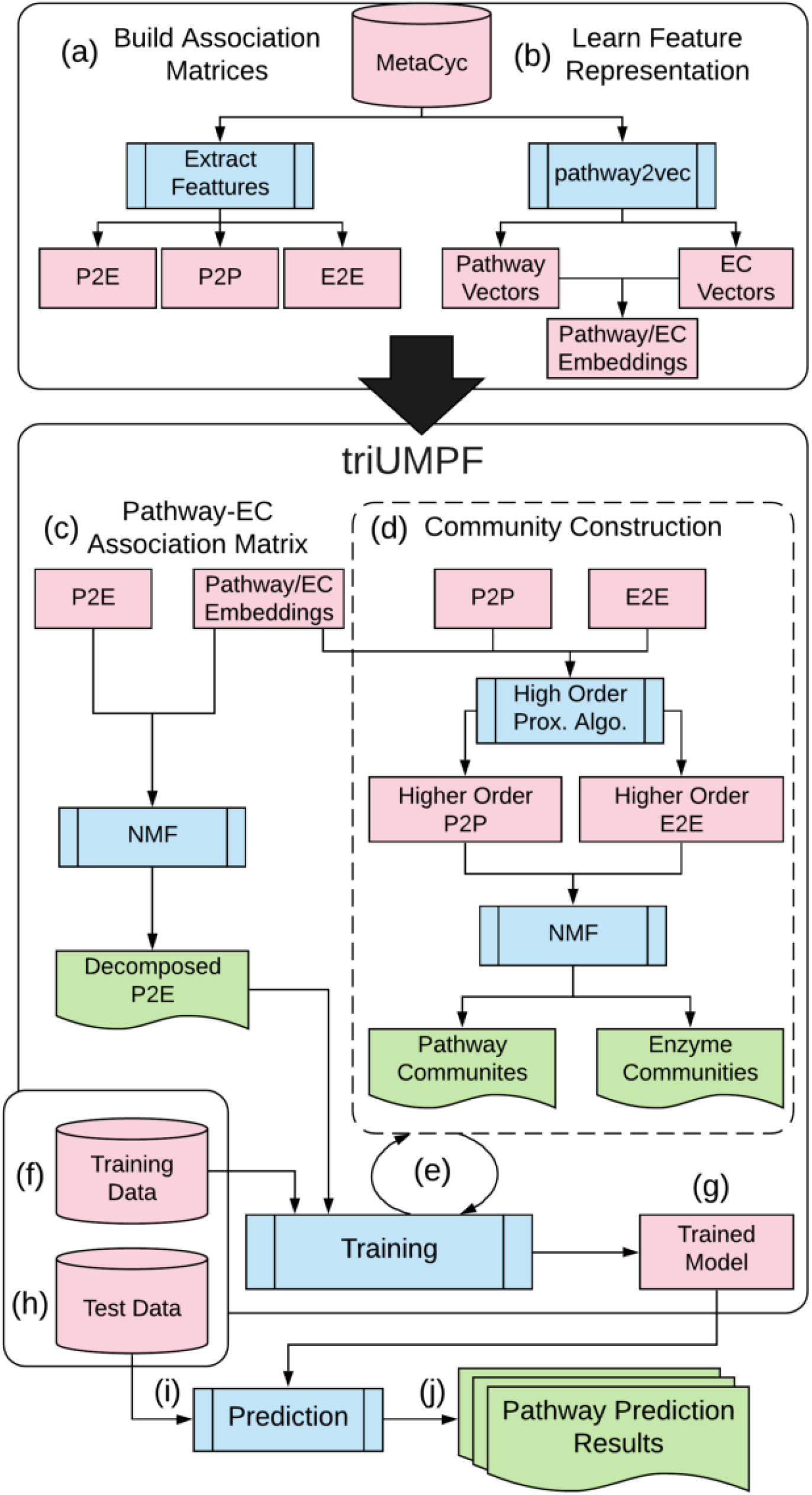
A workflow diagram showing the proposed triUMPF method. Initially, triUMPF takes the Pathway-EC association (P2E) information (a) to produce several low rank matrices (c) while, simultaneously, detecting pathway and EC communities (d) given two interaction matrices, corresponding Pathway-Pathway (P2P) and EC-EC (E2E) (a). For both steps (c) and (d), pathway and EC features obtained from pathway2vec package (b) are utilized. Afterwards, triUMPF iterates between updating community parameters (d) and optimizing multi-label parameters (e) with the use of training data (f). Once the training is achieved the learned model (g) can be used to predict a set of pathways (i-j) from an organismal genome or multi-organismal dataset (h).

### 2.1 Decomposing the Pathway EC Association Matrix

Inspired by the idea of non-negative matrix factorization (NMF), we decompose the P2E association matrix to recover low-dimensional latent factor matrices [9]. Unlike previous application of NMF to biological data [28], triUMPF incorporates constraints into the matrix decomposition process. Formally, let 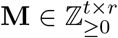 be a non-negative matrix, where *t* is the number of pathways and *r* is the number of enzymatic reactions. Each row in **M** corresponds to a pathway and each column represent an EC, such that **M**_*i,j*_ = 1 if an EC *j* is in pathway *i* and 0 otherwise. Given **M**, the standard NMF decomposes this matrix into the two low-rank matrices, i.e. **M** ≈ **WH**^⊤^, where **W** ∈ ℝ^*t*×*k*^ stores the latent factors for pathways while **H** ∈ ℝ^*r*×*k*^ is latent factors associated with ECs and 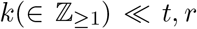. However, triUMPF extends this standard NMF by leveraging features, obtained from *pathway2vec* [24], encoding two interactions: i)- within ECs or pathways and ii)- between pathways and ECs. For more details about this step, please see Appx. Section 5.2.1.

### 2.2 Community Reconstruction and Multi-label Learning

The community detection problem [23,30] is the task of discovering distinct groups of nodes that are densely connected. During this phase, triUMPF performs community detection to guide the learning process for pathways using binary P2P 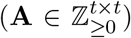 and E2E 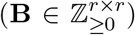 association matrices, where each entry in these matrices is a binary value indicating an interaction among corresponding entities. However, **A** and **B** capture pairwise first-order proximity among their related entities, consequently, they are inadequate to fully characterize distant relationships among pathways or ECs [30]. Therefore, triUMPF utilizes higher-order proximity using the following formula [23]:

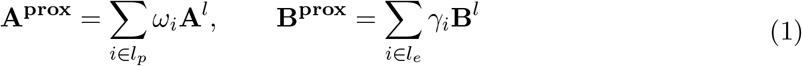

where **A^prox^** and **B^prox^** are polynomials of order *l_p_* ∈ **Z**_>0_ and *l_e_* ∈ **Z**_>0_, respectively, and *ω* ∈ **R**_>0_ and *γ* ∈ **R**_>0_ are weights associated to each term. Using these higher order matrices, triUMPF applies two NMFs to recover communities (Appx. Section 5.2.2). Then, triUMPF uses **W** and **H** from the decomposition phases (Section 2.1) and the detected communities to optimize multi-label pathway parameters in an iterative process (Appx. Section 5.2.3) until the maximum number of allowed iterations is reached. At the end, the trained model can be used to perform pathway prediction from an organismal genome or multi-organismal dataset with high precision due to constraints embedded in the P2E, P2P, and E2E associations matrices.

## 3 Results

We evaluated triUMPF performance across multiple datasets spanning the genomic information hierarchy [25]: i)- T1 golden consisting of EcoCyc, HumanCyc, AraCyc, YeastCyc, LeishCyc, and TrypanoCyc; ii)- three *E. coli* genomes composed of E. coli K-12 substr. MG1655 (TAX-511145), uropathogenic E. coli str. CFT073 (TAX-199310), and enterohemorrhagic E. coli O157:H7 str. EDL933 (TAX-155864); iii)- BioCyc (v20.5 T2 & 3) [5] composed of 9255 PGDBs (Pathway/Genome Databases) with 1463 pathways constructed using Pathway Tools v21 [16]; iv)- *symbionts* genomes of *Moranella* (GenBank NC-015735) and *Tremblaya* (GenBank NC-015736) encoding distributed metabolic pathways for amino acid biosynthesis [26]; v)- Critical Assessment of Metagenome Interpretation (CAMI) initiative low complexity dataset consisting of 40 genomes [31]; and vi)- whole genome shotgun sequences from the Hawaii Ocean Time Series (HOTS) at 25m, 75m, 110m (sunlit) and 500m (dark) ocean depth intervals [33]. We applied BioCyc v20.5 to train triUMPF while the remaining datasets were used to report performance results. Since BioCyc v20.5 contains less than 1460 trainable pathways, we applied pathway2vec with RUST-norm (or “crt”) configuration to improve prediction (see Section 5.4.3). For general statistics about these datasets are summarized in Appx. Table 4.

For comparative analysis, triUMPF’s performance on T1 golden datasets was compared to three pathway prediction methods: i)- MinPath version 1.2 [38], which uses integer programming to recover a conserved set of pathways from a list of enzymatic reactions; ii)- PathoLogic version 21 [16], which is a symbolic approach that uses a set of manually curated rules to predict pathways; and iii)- mlLGPR which uses supervised multi-label classification and rich feature information to predict pathways from a list of enzymatic reactions [25]. In addition to testing on T1 golden datasets, triUMPF performance was compared to PathoLogic on three *E. coli* genomes and to PathoLogic and mlLGPR on mealybug symbionts, CAMI low complexity, and HOTS multi-organismal datasets. The following metrics were used to report on performance of pathway prediction algorithms including: *average precision*, *average recall*, *average F1 score (F1)*, and *Hamming loss* as described in [25]. For experimental settings and additional tests, see Appx. Sections 5.4 and 5.5.

### 3.1 T1 Golden Data

As shown in Table 1, triUMPF achieved competitive performance against the other methods in terms of average precision with optimal performance on EcoCyc (0.8662). However, with respect to average F1 scores, it under-performed on HumanCyc and AraCyc, yielding average F1 scores of 0.4703 and 0.4775, respectively (Appx. Table 5). Since the observed number of pathway labels in BioCyc v20.5 is 1463 pathways (a subset of 2526 MetaCyc pathways) (as explained in Section 3), triUMPF trained with this data (using features from pathway2vec [24]) can not infer pathways outside the trainable pathways. Consequently, this has translated into low average F1 scores of HumanCyc and AraCyc. A possible treatment would be incorporating additional PGDBs containing more pathways to train triUMPF. However, this would require substantially building many PGDBs from organismal genomes or using multiple versions of BioCyc data. A detailed analysis on this is left for future work.

**Table 1:**
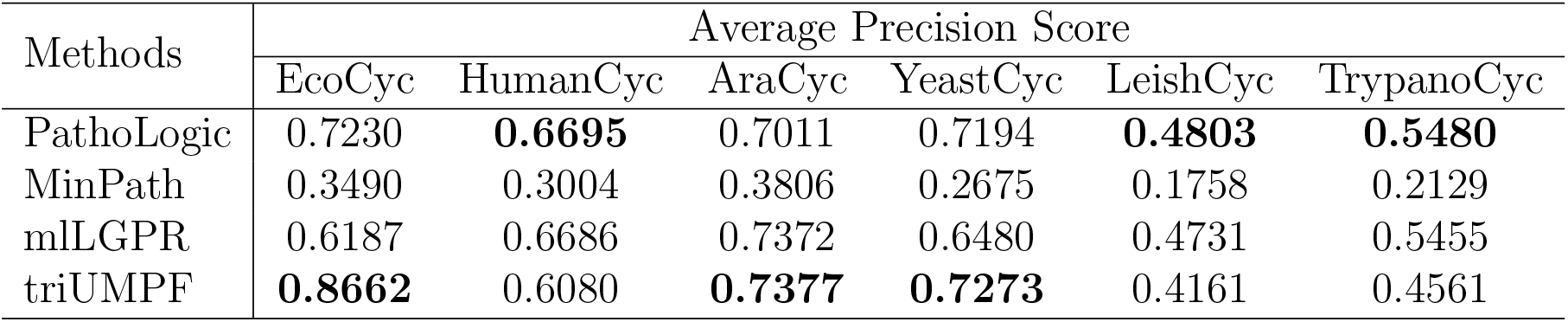
Average precision of each comparing algorithm on 6 golden T1 data.

### 3.2 Three E. coli data

Recall that community detection (Section 2.2) was used to guide the multi-label learning process. To demonstrate the influence of communities on pathway prediction, we compared pathways predicted for the T1 gold standard E. coli K-12 substr. MG1655 (TAX-511145), henceforth referred to as MG1655, using PathoLogic and triUMPF. Appx. Fig. 8a shows the results, where both methods inferred 202 true-positive pathways (green-colored) in common out of 307 expected true-positive pathways (using EcoCyc as a common frame of reference). In addition, PathoLogic uniquely predicted 39 (magenta-colored) true-positive pathways while triUMPF uniquely predicted 16 true-positives (purple-colored). This difference arises from the use of tax-onomic pruning in PathoLogic which improves recovery of taxonomically constrained pathways and limits false-positive identification. With taxonomic pruning enabled, PathoLogic inferred 79 false-positive pathways, and over 170 when pruning was disabled. In contrast triUMPF which does not use taxonomic feature information inferred 84 false-positive pathways (Appx. Table 6). This improvement over PathoLogic with pruning disabled reinforces the idea that pathway communities improve precision of pathway prediction with limited impact on overall recall. Based on these results, it is conceivable to train triUMPF on subsets of organismal genomes resulting in more constrained pathway communities for pangenome analysis.

To further evaluate triUMPF performance on closely related organismal genomes, we performed pathway prediction on E. coli str. CFT073 (TAX-199310), and E. coli O157:H7 str. EDL933 (TAX-155864) and compared results to the MG1655 reference strain [36]. Both CFT073 and EDL933 are pathogens infecting the human urinary and gastrointestinal tracts, respectively. Previously, Welch and colleagues described extensive genomic mosaicism between these strains and MG1655, defining a core backbone of conserved metabolic genes interspersed with genomic islands encoding common pathogenic or niche defining traits [36]. Neither CFT073 nor EDL933 genomes are represented in the BioCyc collection of organismal pathway genome databases. A total of 335 and 319 unique pathways were predicted by PathoLogic and triUMPF, respectively. The resulting pathway lists were used to perform a set-difference analysis with MG1655 (Fig. 2). Both methods predicted more than 200 pathways encoded by all three strains including core pathways like the *TCA* cycle (Appx. Figs 8b and 8c). CFT073 and EDL933 were predicted to share a single common pathway (*TCA cycle IV (2-oxoglutarate decarboxylase)*) by triUMPF. However this pathway variant has not been previously identified in E. coli and is likely a false-positive prediction based on recognized taxonomic range. Both PathoLogic and triUMPF predicted the *aerobactin biosynthesis* pathway involved in siderophore production in CFT073 consistent with previous observations [36]. Similarly, four pathways (e.g. *L-isoleucine biosynthesis III* and *GDP-D-perosamine biosynthesis*) unique to EDL933 were inferred by both methods.

**Figure 2:**
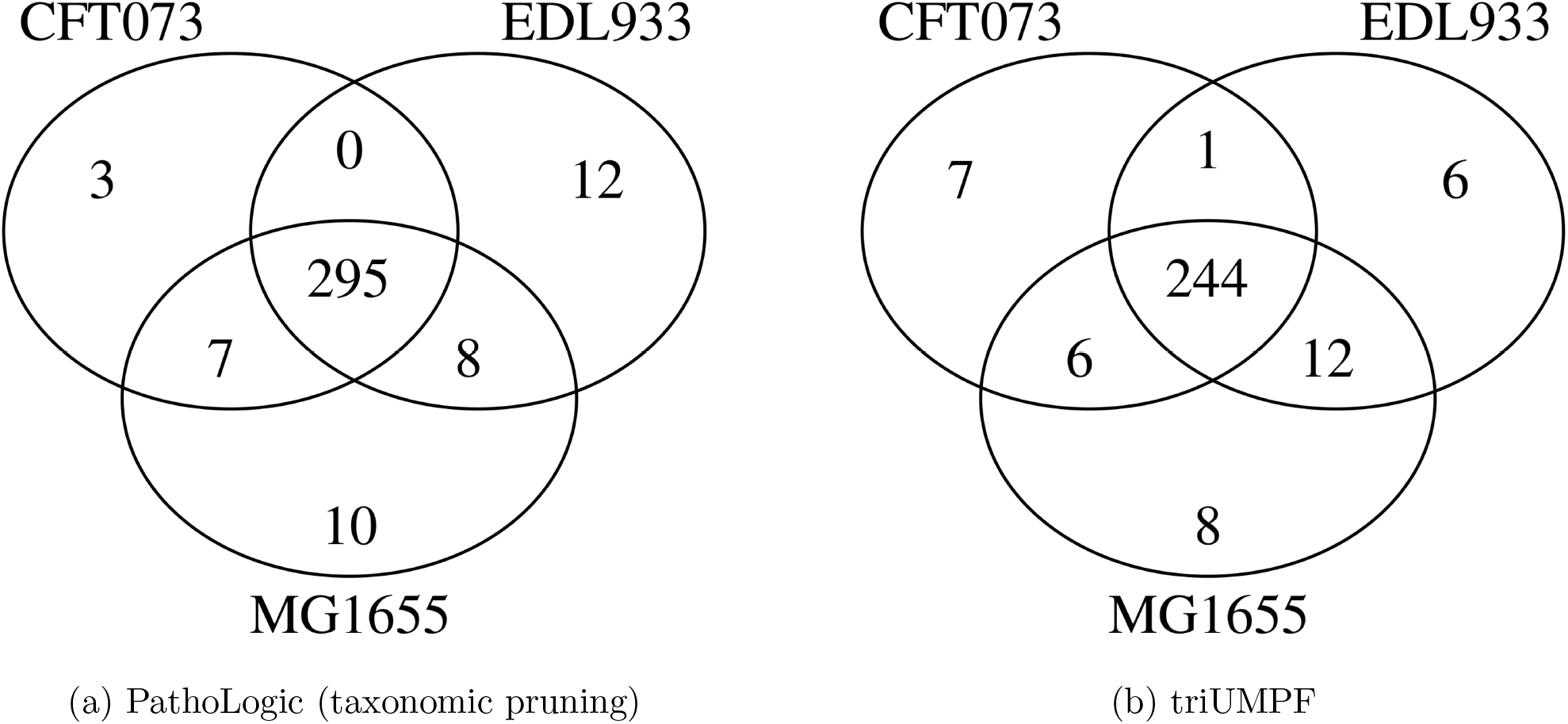
A three way set difference analysis of pathways predicted for E. coli K-12 substr. MG1655 (TAX-511145), E. coli str. CFT073 (TAX-199310), and E. coli O157:H7 str. EDL933 (TAX-155864) using (a) PathoLogic (taxonomic pruning) and (b) triUMPF.

Given the lack of cross validation standards for CFT073 and EDL933 we were unable to determine which method inferred fewer false-positives across the complete set of predicted pathways. To constrain this problem on a subset of the data, we applied GapMind [29] to analyze amino acid biosynthesis pathways encoded in MG1655, CFT073 and EDL933 genomes. GapMind is a web-based application developed for annotating amino acid biosynthesis pathways in prokaryotic microorganisms (bacteria and archaea), where each reconstructed pathway is supported by a confidence level. After excluding pathways that were not incorporated in the training set, a total of 102 pathways were identified across the three strains encompassing 18 amino acid biosynthesis pathways and 27 pathway variants with high confidence (Appx. Table 7). PathoLogic inferred 49 pathways identified across the three strains encompassing 15 amino acid biosynthesis pathways and 17 pathway variants while triUMPF inferred 54 pathways identified across the three strains encompassing 16 amino acid biosynthesis pathways and 19 pathway variants including *L-methionine biosynthesis* in MG1655, CFT073 and EDL933 that was not predicted by PathoLogic. Neither method was able to predict *L-tyrosine biosynthesis I* (Appx. Fig. 10).

### 3.3 Mealybug Symbionts data

To evaluate triUMPF performance on distributed metabolic pathways, we used the reduced genomes of *Moranella* and *Tremblaya* [26]. Collectively the two symbiont genomes encode intact biosynthesis pathways for 9 essential amino acids. PathoLogic, mlLGPR, and triUMPF were used to predict pathways on individual symbiont genomes and a composite genome consisting of both, and resulting amino acid biosynthesis pathway distributions were determined (Fig. 3). Both triUMPF and PathoLogic predicted 6 of the expected amino acid biosynthesis pathways on the composite genome while mlLGPR predicted 8 pathways. The pathway for phenylalanine biosynthesis (*L-phenylalanine biosynthesis I*) was excluded from analysis because the associated genes were reported to be missing during the ORF prediction process. False positives were predicted for individual symbiont genomes in *Moranella* and *Tremblaya* using both methods although pathway coverage was reduced in relation to the composite genome.

**Figure 3:**
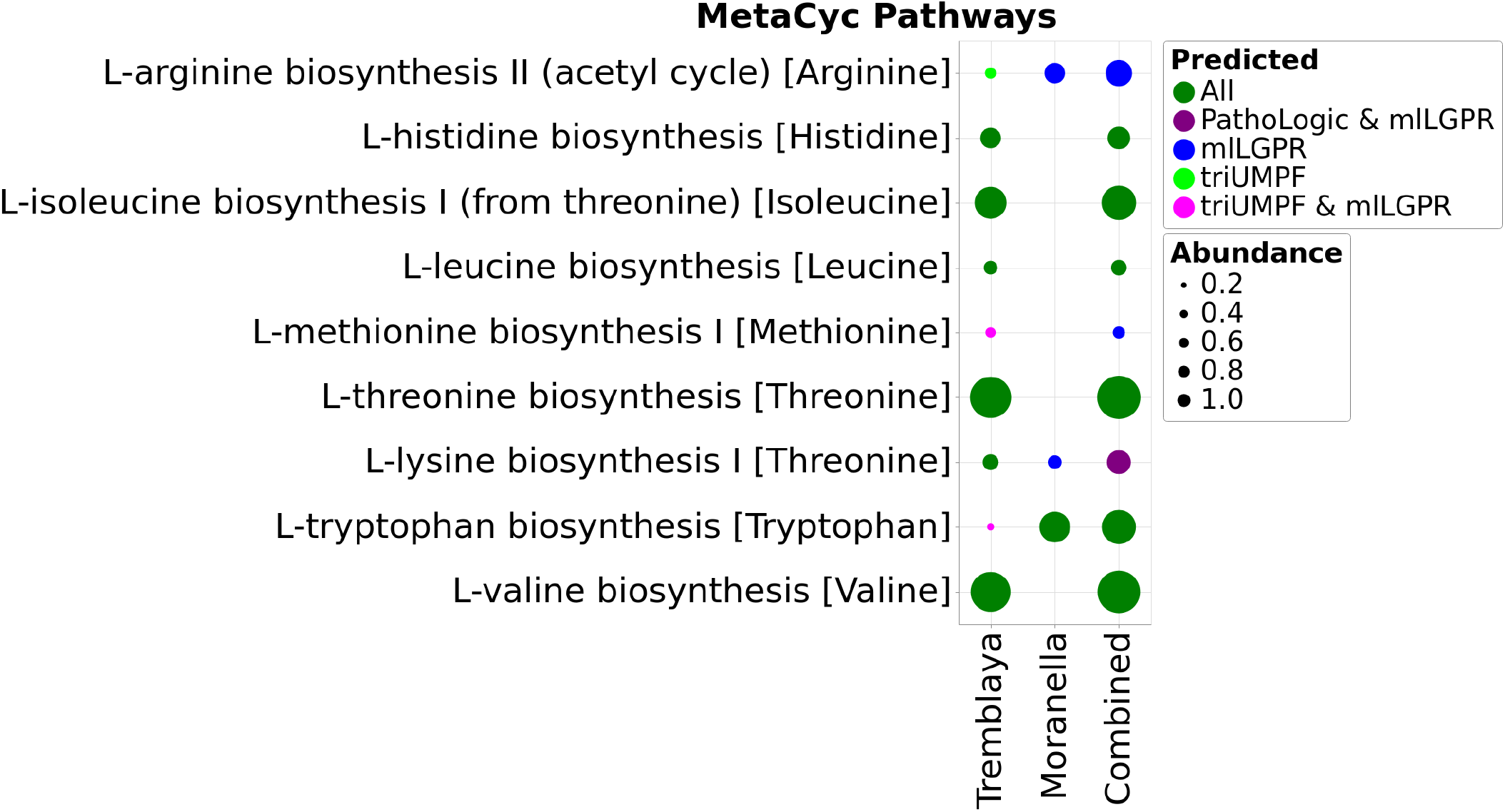
Comparative study of predicted pathways for symbiotic data between PathoLogic, mlL-GPR, and triUMPF. The size of circles corresponds the associated coverage information.

### 3.4 CAMI and HOTS data

To evaluate triUMPF’s performance on more complex multi-organismal genomes, we used the CAMI low complexity [31] and HOTS datasets [33] comparing resulting pathway predictions to both PathoLogic and mlLGPR. For CAMI low complexity, triUMPF achieved an average F1 score of 0.5864 in comparison to 0.4866 for mlLGPR which is trained with more than 2500 labeled pathways (Table 2). Similar results were obtained for HOTS (see Appx. Section 5.5.4).

**Table 2:**
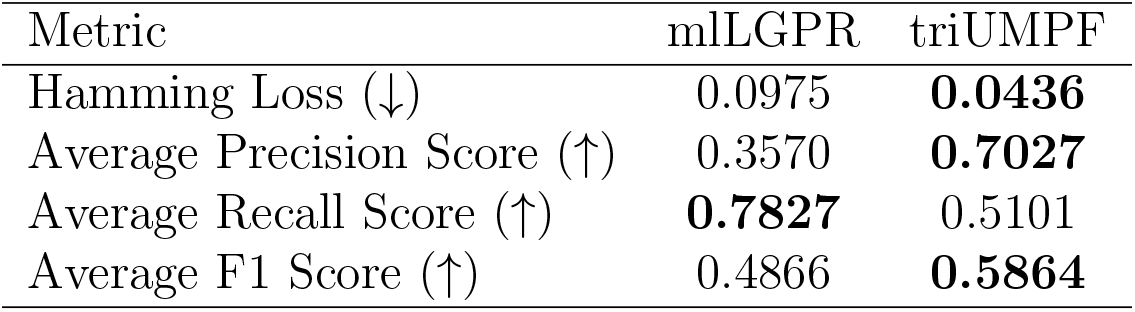
Predictive performance of mlLGPR and triUMPF on CAMI low complexity data. For each performance metric, ‘↓’ indicates the smaller score is better while ‘↑’ indicates the higher score is better.

Among a subset of 180 selected water column pathways, PathoLogic and triUMPF predicted a total of 54 and 58 pathways, respectively, while mlLGPR inferred 62. From a real world perspective none of the methods predicted pathways for *photosynthesis light reaction* nor *pyruvate fermentation to (S)-acetoin* although both are expected to be prevalent in the water column. Perhaps, the absence of specific ECs associated with these pathway limits rule-based or ML prediction. Indeed, closer inspection revealed that the enzyme *catabolic acetolactate synthase* was missing from the *pyruvate fermentation to (S)-acetoin* pathway, which is an essential rule encoded in PathoLogic and represented as a feature in mlLGPR. Conversely, although this pathway was indexed to a community, triUMPF did not predict its presence, constituting a false-negative.

## 4 Discussion and Conclusion

In this paper we introduced a novel ML approach for metabolic pathway inference that combines three stages of NMF to capture relationships between enzymes and pathways within a network followed by community detection to extract higher order network structure. First, a Pathway-EC association (**M**) matrix, obtained from MetaCyc, is decomposed using the NMF technique to learn a constrained form of the pathway and EC factors, capturing the microscopic structure of **M**. Then, we obtain the community structure (or mesoscopic structure) jointly from both the input datasets and two interaction matrices, Pathway-Pathway interaction and EC-EC interaction. Finally, the consensus relationships between the community structure and data, and between the learned factors from **M** and the pathway labels coefficients are exploited to efficiently optimize metabolic pathway parameters.

We evaluated triUMPF performance using a corpora of experimental datasets manifesting diverse multi-label properties comparing pathway prediction outcomes to other prediction methods including PathoLogic [16] and mlLGPR [25]. During benchmarking we realized that the BioCyc collection suffers from a class imbalance problem [13] where some pathways infrequently occur across PGDBs. This results in a significant sensitivity loss on T1 golden data, where triUMPF tended to predict more frequently observed pathways while missing more infrequent pathways. One potential approach to solve this class-imbalance problem is subsampling the most informative PGDBs for training, hence, reducing false-positives [19]. Despite the observed class imbalance problem, triUMPF improved pathway prediction precision without the need for taxonomic rules or EC features to constrain metabolic potential. From an ML perspective this is a promising outcome considering that triUMPF was trained on a reduced number of pathways relative to mlLGPR. Future development efforts will explore subsampling approaches to improve sensitivity and the use of constrained taxonomic groups for pangenome and multi-organismal genome pathway inference.

## 5 Appendices

## 5.1 Appendix A1: Definitions and Problem Formulation

Here, the default vector is considered to be a column vector and is represented by a boldface lowercase letter (e.g., **x**) while matrices are represented by boldface uppercase letters (e.g., **X**). The **X**_*i*_ matrix indicates the *i*-th row of **X** and **X**_*i,j*_ denotes the (*i, j*)-th entry of **X** while, for a vector, **x**_*i*_ denotes an *i*-th cell of **x**. The transpose of **X** is denoted as **X**^⊤^ and the trace of it is symbolized as **tr**(**X**). The Frobenius norm of **X** is defined as 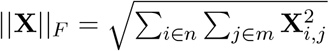. Occasional superscript, **x**^(*i*)^, suggests an index to a sample, a power, or a current epoch during a learning period. We use calligraphic letters to represent sets (e.g., *ε*) while we use the notation |.| to denote the cardinality of a given set. With these notations in mind, we introduce several concepts integral to the problem formulation.

Metabolic pathway inference from genomic sequence information at different levels of complexity and completion requires a trusted source of labeled pathway information in which the set of ordered reactions within and between cells is linked to substrates and products (compounds or metabolites). This information can be represented in graphs corresponding to reactome and pathway-level interactions. In this study, we use MetaCyc, a multi-organism member of the BioCyc collection of Pathway/Genome Databases (PGDB) as the trusted source for reactome and pathway information [5]. MetaCyc contains only experimentally validated metabolic pathways across all domains of life. To simplify computational complexity, we consider the reaction and pathway graphs to be undirected.

### Definition 5.1. Reaction Graph Topology.

Let the reaction graph be represented by an undirected graph 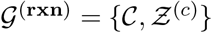, where 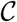 is a set of *c* metabolites and 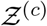 represents *r*′ links between compounds. Each link indicates a reaction, derived from a set of biochemical reactions 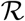 of size *r*′. Then, the reaction graph topology is defined by a matrix 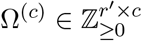, where each entry 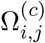 is a binary value of 1 or 0, indicating either the compound *j* is a substrate/product in a reaction *i* or not involved in that reaction, respectively.

### Definition 5.2. Pathway Graph Topology.

Let 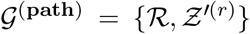 be an undirected graph, where 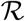 is presented in Def. 5.1, and 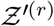 represents a set of *t*′ links between reactions. Then, the pathway graph topology is defined by a matrix 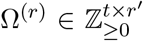, where each entry 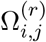 is either 0 or a positive integer, corresponding the absence or the frequency of the reaction *j* in pathway *i*, respectively. And, *t* is the number of pathways in a set 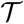.

Note that reactions in 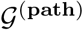 may be annotated as a *spontaneous reaction* or a reaction catalyzed by one or more enzymes, *enzymatic reaction* and classified by an *enzyme commission* number (EC) [27]. In addition, a number of enzymes referred to as *promiscuous enzymes* can participate in more than one pathway. Given this information we associate EC numbers to pathways and formulate three graphs, one representing associations between pathways and enzymes indicated by enzyme commission (EC)) numbers, one representing interactions between enzymes and another representing interactions between pathways.

### Definition 5.3. Pathway-EC Association (P2E).

Let 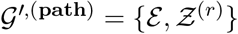 be a subgraph of 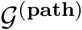, such that 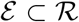 with *r* ≪ *r*′ enzymatic reactions. Then, the Pathway-EC association is defined as a matrix 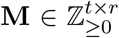, where each row corresponds to a pathway, and each column represent an EC, such that **M**_*i,j*_ = 1 if an EC *j* is in pathway *i* and 0 otherwise.▪

Typically, the association matrix **M** is extremely sparse. Using reaction and pathway graph topology, we build interaction adjacency matrices as follows.

### Definition 5.4. EC-EC Interaction (E2E).

Given 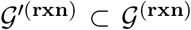 we define an EC-EC interaction matrix 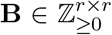 such that an entry **B**_*i,j*_ is a binary value encoding an interaction between two ECs *i* and *j* iff they both share a compound, i.e., 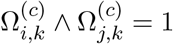 where 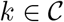.▪

### Definition 5.5. Pathway-Pathway Interaction (P2P).

Given 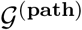, we define a Pathway-Pathway interaction matrix 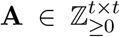 such that an entry **A**_*i,j*_ is a binary value indicating an interaction between pathways *i* and *j* if there exists a reaction 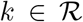 where associated compounds are either substrate or product in both *i* and *j* pathways.▪

After determining relationships within each graph, we define a *multi-label* metabolic pathway dataset.

### Definition 5.6. Multi-label Pathway Dataset[25].

A general form of pathway dataset is characterized by 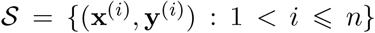 consisting of *n* examples, where **x**^(*i*)^ is a vector indicating the abundance information corresponding to each enzymatic reaction. An enzymatic reaction, in turn, is denoted by *e*, which is an element of a set of enzymatic reactions *ε* = {*e*_1_*, e*_2_*,…, e_r_*}, having *r* possible reactions. The abundance of an enzymatic reaction *i*, for example 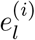, is defined as 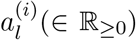. The class labels 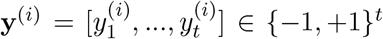 is a pathway label vector of size *t* that represents the total number of pathways, which are derived from a set of labeled metabolic pathway 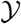. The matrix form of **x**^(*i*)^ and **y**^(*i*)^ are symbolized as **X** and **Y**, respectively.▪

The input space is assumed to be encoded as *r*-dimensional feature vector and is symbolized as 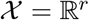. Furthermore, each example in 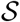 is considered to be drawn independent, identically distributed (i.i.d) from an unknown distribution 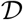 over 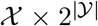. Now we state the problem considered in this paper.

## Metabolic Pathway Prediction.

Given: i)- Pathway-EC matrix **M**, ii)- a Pathway-Pathway interaction matrix **A**, iii)- an EC-EC interaction matrix **B**, and iv)- a dataset 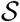, the goal is to efficiently reconstruct pathway labels for a hitherto unseen instance **x**^∗^.

## 5.2 Appendix A2: Detailed Description of triUMPF Method

In this section, we provide a description of triUMPF components, presented in Fig. 1 of main manuscript, including: i)- decomposing the pathway EC association matrix, ii)- subnetwork or community reconstruction, and iii)- the multi-label learning process.

## 5.2.1 Decomposing the Pathway EC Association Matrix

Given the non-negative **M**, we formulate the following minimization objective function:

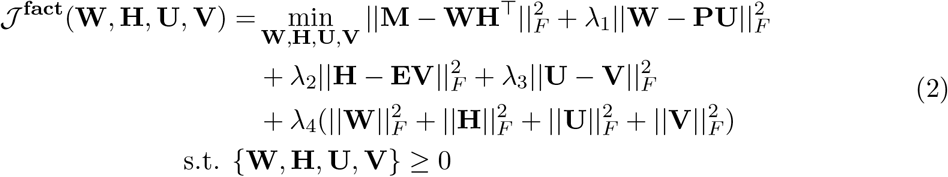

where **W** ∈ ℝ^*t*×*k*^ stores the latent factors for pathways while **H** ∈ ℝ^*r*×*k*^, known as the basis matrix, can be thought of as latent factors associated with ECs and *k* ≪ *t, r* and *λ*_∗_ are regularization hyperparameters. The leftmost term is the well-known squared loss function that penalizes the deviation of the estimated entries in both **W** and **H** from the true association matrix **M**. The second term corresponds to the relative differences of latent matrix **W** from the pathway features **P** ∈ ℝ^*t*×*m*^, learned using pathway2vec framework, where the matrix **U** ∈ ℝ^*m*×*k*^ absorbs different scales of matrices **W** and **P**. Similarly, the third term indicates the squared loss of **H** from **E** ∈ ℝ^*r*×*m*^, which denotes the feature matrix of ECs, and their differences are captured by **V** ∈ ℝ^*m*×*k*^. In the fourth term, we minimize the differences between factors **U** and **V**, capturing shared prominent features for the low dimensional coefficients.

## 5.2.2 Subnetwork or Community Reconstruction

Recall from the main manuscript, the higher order proximity of the two matrices **A** and **B** is defined according to the formula [23]:

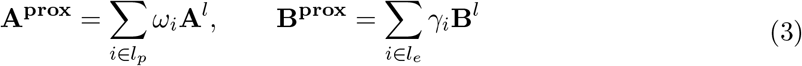

where **A^prox^** and **B^prox^** are polynomials of order *l_p_* ∈ **Z**_>0_ and *l_e_* ∈ **Z**_>0_, respectively, and *ω* ∈ **R**_>0_ and *γ* ∈ **R**_>0_ are weights associated to each term. Using these higher order matrices, we invoke NMF to recover communities.

Formally, let **T** ∈ ℝ^*m*×*p*^ be a non-negative community representation matrix of size *p* communities for pathways, where the *j*-th column in **T**_:*,j*_ denotes the representation of community *j*. The pathway community indicator matrix is denoted by **C** ∈ ℝ^*t*×*p*^ conditioned on **tr**(**C**^⊤^**C**) = *t*, where each entry **C**_*i,l*_ and **C**_*j,l*_ encodes the probability that pathways *i* and *j* generates an edge belonging to a community *l*. The probability of *i* and *j* belonging to the same community can be assessed as: 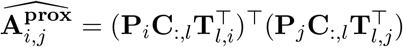. Similar discussion follows for the non-negative representation matrix **R** ∈ ℝ^*m*×*v*^ and the EC community indicator matrix **K** ∈ ℝ^*r*×*v*^ of *v* communities, conditioned on **tr**(**K**^⊤^ **K**) = *r*. Unfortunately, due to the constraints emphasized on **C** and **K**, it is not straightforward to analytically derive an expression, instead, we resort to a more tractable solution provided in [35], and relax the condition to be an orthogonal constraint, resulting in the following objective function:

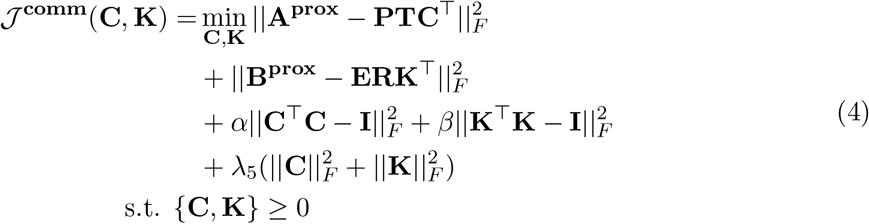

where **I** denotes an identify matrix, *λ*_5_ is a regularization hyperparameter while *α* and *β* are both positive hyperparameters. The value of these hyperparameters is usually set to a large number, e.g. 10^9^ in this work, for adjusting the contribution of corresponding terms. The obtained communities in Eq 4 are directly linked to the underlying graph topologies, i.e., **A^prox^** and **B^prox^**.

## 5.2.3 Multi-label Learning Process

We now bring together the NMF and community detection steps with multi-label classification for pathway prediction. The learning problem must balance between information in **M** while being lenient towards the dataset 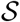, which should provide enough evidence to generate representations of communities among pathways and ECs, as suggested by **A^prox^** and **B^prox^**. We present a weight term Θ ∈ ℝ^*t*×*r*^ that enforces **X** to be close enough to both **Y** and **M**. We also introduce two auxiliary terms **L** ∈ ℝ^*n*×*m*^, which capture correlations between **X** and **Y** and **Z** ∈ ℝ^*r*×*r*^, enforcing the pathway coefficients associated with **M** resulting in the following objective function:

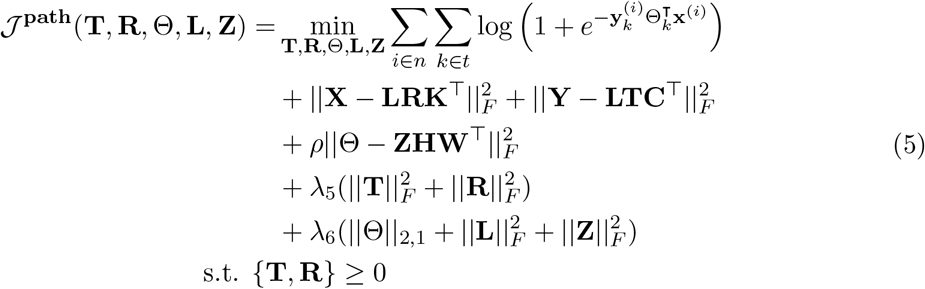

where *λ*_5_, *λ*_6_, and *ρ* are regularization hyperparameters, and ||.||_2,1_ represents the sum of the Euclidean norms of columns of a matrix introduced to emphasize sparseness. Notice that we do not restrict the terms **L** and **Z** to be non-negative. Both the second and the third terms in Eq. 5, are needed to discover pathway and EC communities, i.e., **C** and **K**, respectively.

The Eqs 2, 4, and 5 are jointly non-convex due to non-negative constraints on the original and the approximation factorized matrices, implying the solutions to triUMPF are only unique up to scalings and rotations [37]. Hence, we adopt an alternating optimization algorithm to solve each objective function simultaneously, which is provided in Section 5.3.

## 5.3 Appendix A3: Optimization

In this section, we derive the optimization for triUMPF’s objective function:

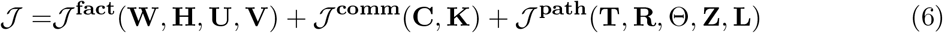

where,

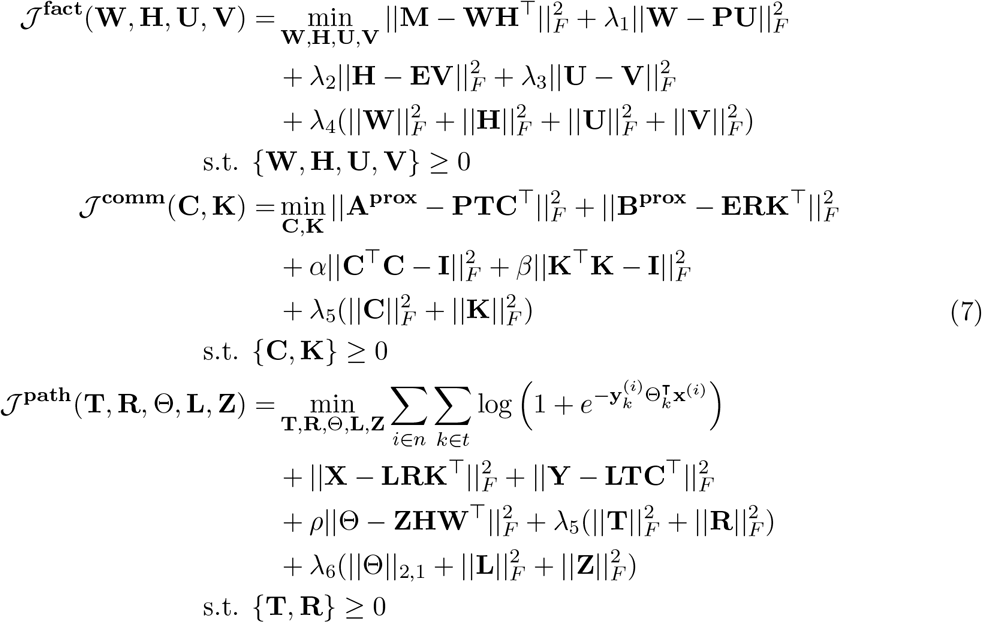

The objective function in Eq. 7 is non-convex due to multiple non-negative constraints. Numerous algorithms have been proposed to optimize the objective function, including alternating non-negative least squares [17] and hierarchical alternating least squares [6]. Here, we employ the original algorithm for NMF which was introduced in [22] and consists of simple multiplicative update rules (with auxiliary variables) that are based on the gradient descent technique [10]. Beginning with random positive initialization, element-wise updates of Eq 6 w.r.t **W**, **H**, **U**, **V**, **C**, **K**, **T**, **R**, Θ, **Z**, and **L** at each iteration are applied until convergence. The gradient descent aims to search for a local minima of the cost function by moving in the direction of its steepest descent. By introducing Lagrangian multipliers (auxiliary variables), which are *ψ*, *ϕ*, *φ*, *ϱ*, *ζ*, *ϖ*, *κ*, and *ξ* to enforce the constraints for **W**, **H**, **U**, **V**, **C**, **T**, **R**, **K**, respectively, Eq. 7 can be reformulated as:

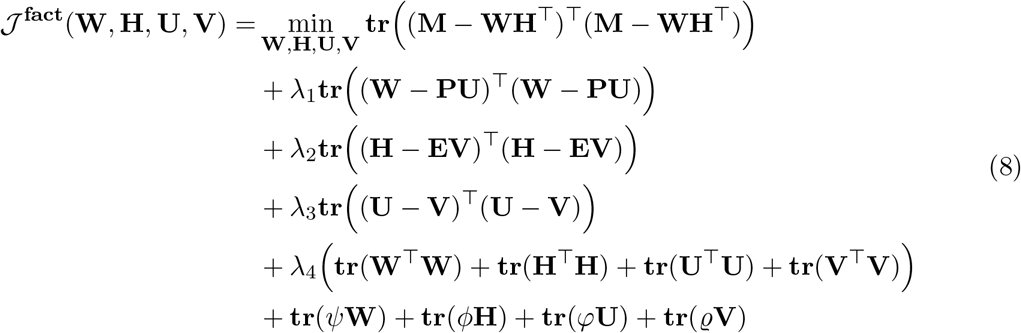

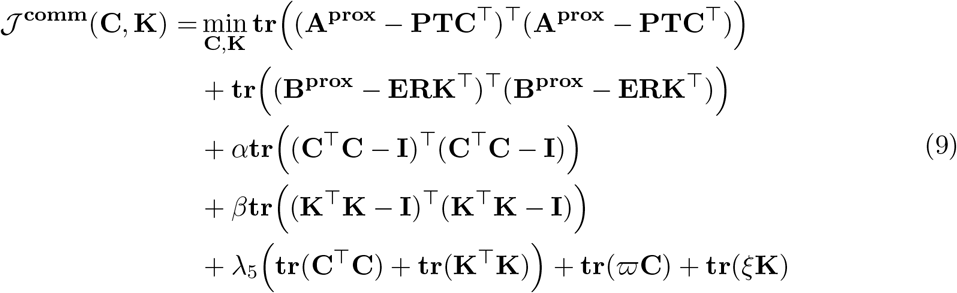

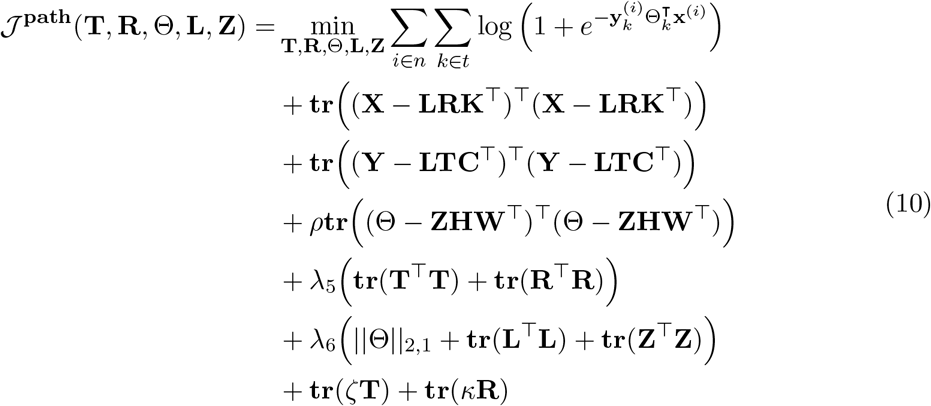

where **tr**(.) denotes the trace of a matrix. Using the addition property of the transpose, (**X**+ **Y**)^⊤^ = **X**^⊤^ + **Y**^⊤^, and its multiplication property, (**XY**)^⊤^ = **Y**^⊤^**X**^⊤^, we can expand the trace of the first term as

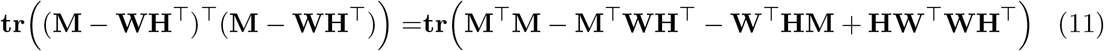

By expanding the remaining terms in Eq. 8 and using the trace of a sum of matrix property, **tr**(**X**+ **Y**) = **tr**(**X**) + **tr**(**Y**), we obtain the following formula:

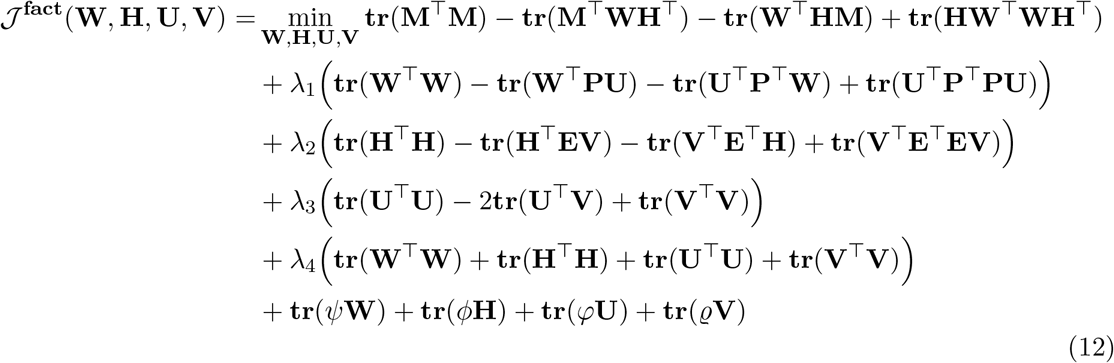

Similar to the process of getting Eq. 12, we expand the Eq. 9 as:

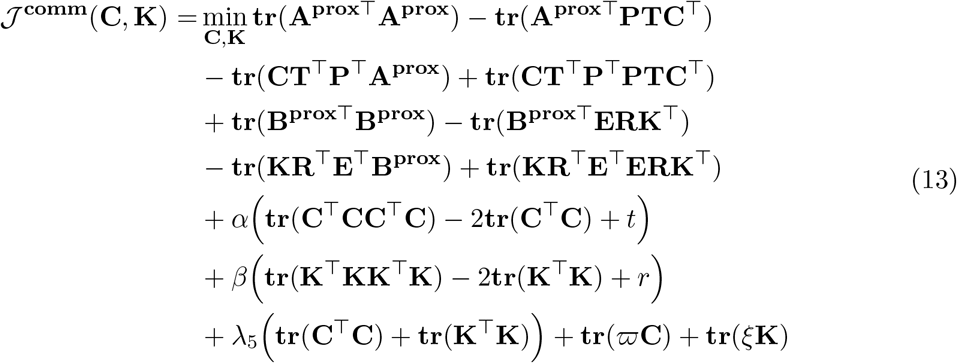

Expand the Eq. 10, we obtain the following:

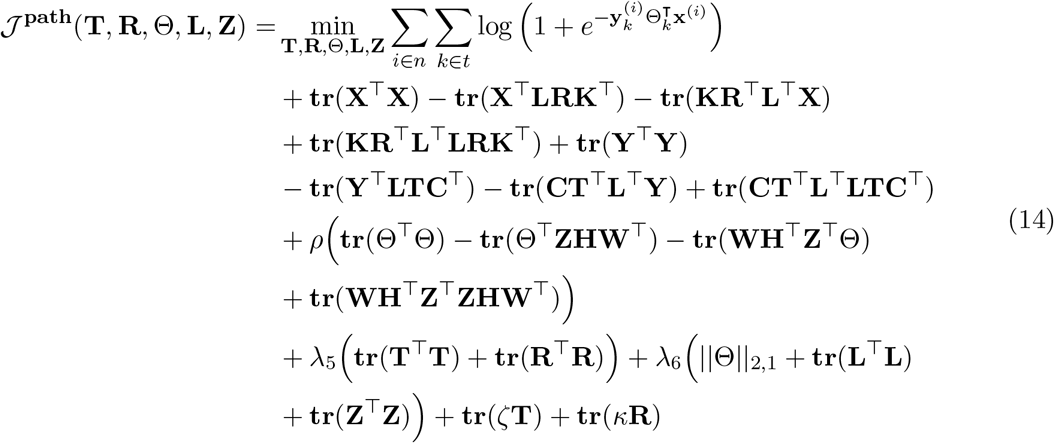

As explained earlier, the objective functions in Eqs 12, 13, and 14, are not convex with respect to all parameters combined. Instead in NMF, **W**, **H**, **U**, **V**, **C**, **K**, **T**, **R**, Θ, **L**, and **Z** are individually optimized in an iterative process, where we update one matrix at a time while keeping the remaining matrices fixed. This ensures convergence to a local minima for each subproblem. This methods is called *block-coordinate descent*. Hence, the update of parameters occur in the following four alternate optimization steps for 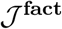: i)- the basis matrix **W**, representing pathway factors, ii)- the latent coefficient matrix **H**, representing EC factors, iii)- the linear transformation **U**, and iv)- the other linear transformation **V**. For 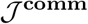, we alternate between the community indicator matrix **C** for pathways and the other community indicator matrix **K** for ECs. Finally, we optimize, alternatively, the two community representation matrices **T** and **R** for pathways and ECs, respectively, the two auxiliary matrices **L** and **Z**, and the input weight matrix Θ. The three objective functions, 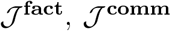, and 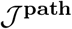 are run simultaneously in a divide and conquer strategy. Detailed rules for updating all the variables are outlined below.

## 1. Update the basis matrix W.

To update the feature matrix **W**, we fix **H**, **U** and **V**. Then, the objective function in Eq. 12 w.r.t **W** is reduced to the following formula (after dropping the min operation):

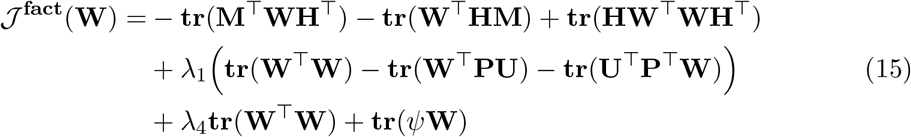

where *ψ* is the Lagrange multiplier for the constraint **W** ≥ 0. For computing the gradient of this equation, we use the following properties with respect to **X**:

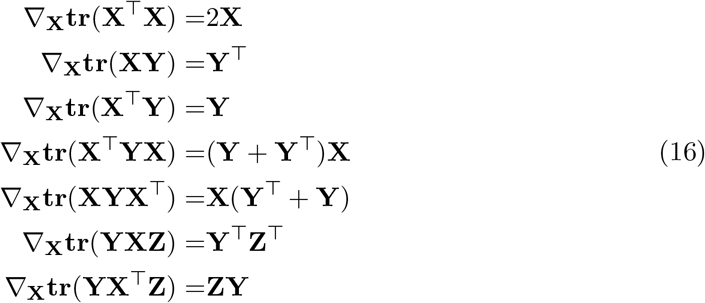

By computing the gradient of the cost function in Eq. 15 w.r.t **W** to 0, we have:

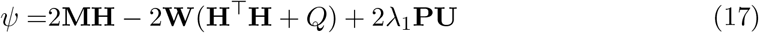

where *Q* = (*λ*_1_ + *λ*_4_). Following the Karush-Kuhn-Tucker (KKT) condition for the non-negativity of **W**, we have the following equation:

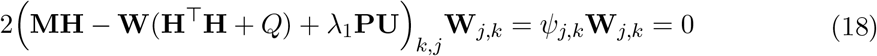

Given an initial value of **W**, the successive updating rule of **W** is:

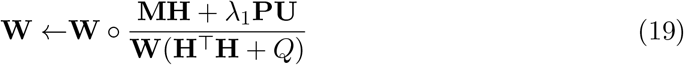

The iterative update rules in Eq. 19 is transformed into multiplicative update rules, which cannot generate negative elements since all values are positive and only multiplications and divisions are involved at each iteration [21].

## 2. Update the latent coefficient matrix H.

The feature matrix **H** is updates as described above in which **W**, **U** and **V** are fixed to obtain the objective function for Eq. 12 w.r.t **H** as:

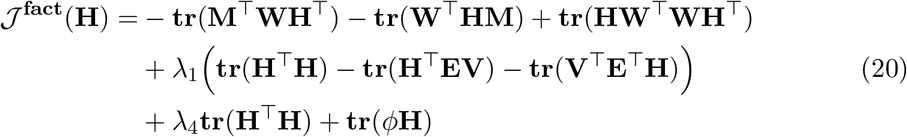

Taking the derivative of the cost function in Eq. 20 w.r.t **H** to 0 and using the gradient properties in Eq. 16, we obtain the following:

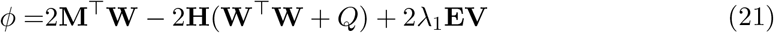

where *Q* = (*λ*_1_ + *λ*_4_). With the KKT complementary condition for the nonnegativity of **H**, we have:

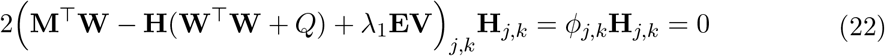

The multiplicative updates after some algebraic manipulation w.r.t parameter **H**:

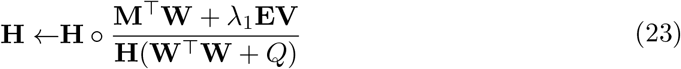

## 3. Update the linear transformation U.

Suppose that **W**, **H** and **V** are fixed, then Eq. 12 w.r.t **U** is reduced to:

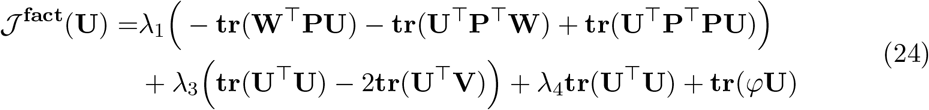

Then we take the derivative of above formula with respect to the transformation matrix **U** to 0:

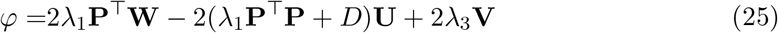

where *D* = (*λ*_3_ + *λ*_4_). Formulating the above equation based on Karush–Kuhn–Tucker conditions for the nonnegativity of **U** results in:

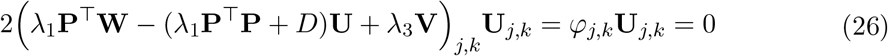

Then, the parameter **U** is updated according to:

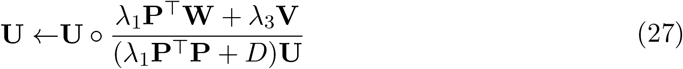

## 4. Update the linear transformation V.

To update the linear transformation matrix **V**, that **W**, **H** and **U** are fixed, then the transformation matrix **V** is updated such that the error is minimized:

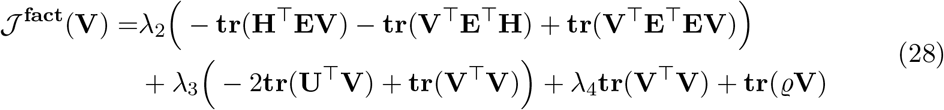

Taking the derivative of this error with respect to **V** to 0 and after some manipulations, we have:

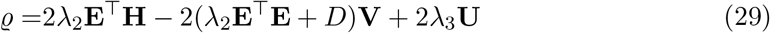

where *D* = (*λ*_3_ +*λ*_4_). Following the Karush–Kuhn–Tucker conditions for the nonnegativity of **V**, we have:

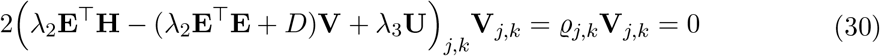

As usual, the parameter **V** is updated according:

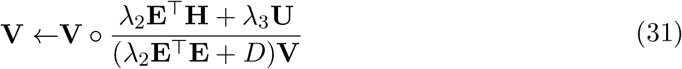

## 5. Update the community indicator matrix C for pathways.

In a similar process, we fix **K**, and update **C**. The matrix **C** is updated such that the error is minimized:

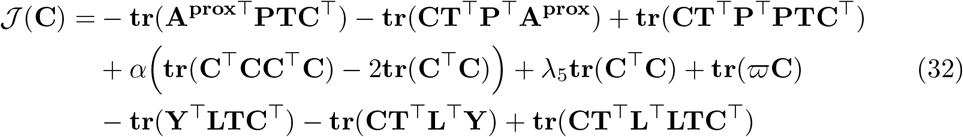

Taking the derivative of this error with respect to **C** to 0, we have:

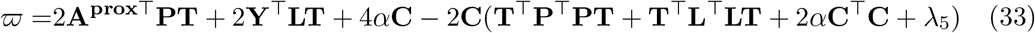

Again, we follow the Karush–Kuhn–Tucker conditions for the nonnegativity of **C**

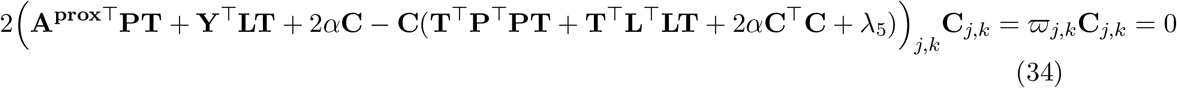

The parameter **C** is updated according:

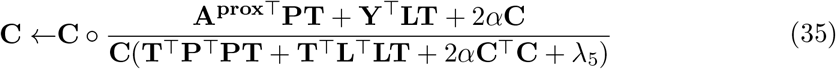

## 6. Update the community indicator matrix K for ECs.

Once the parameter **C** is updated, we use it to update **K**. The matrix **K** is updated such that the error is minimized:

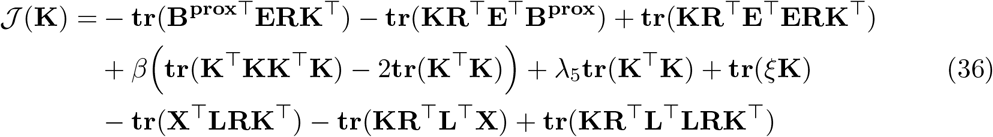

Taking the derivative of this error with respect to **K** to 0, we have:

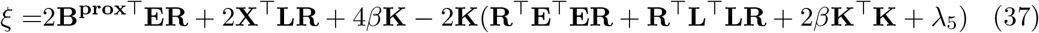

Using the Karush–Kuhn–Tucker conditions for the nonnegativity of **K**, we obtain:

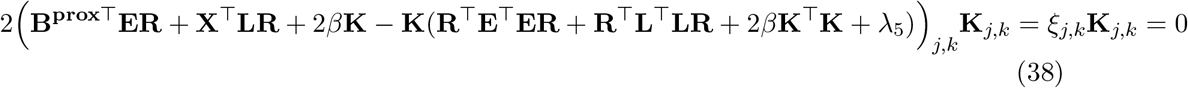

The parameter **K** is updated according:

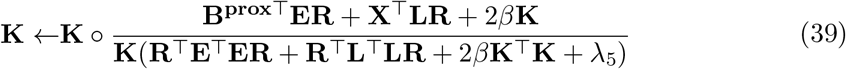

## 7. Update the community representation matrix T for pathways.

By fixing the parameters **C**, **R**, and **K**, we update **T**. The matrix **T** is updated such that the error is minimized:

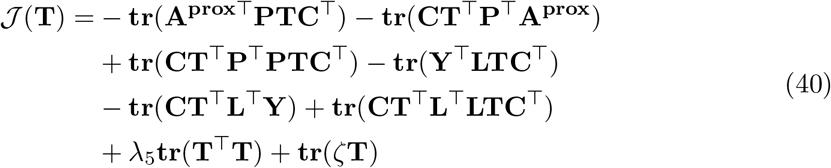

Taking the derivative of this error with respect to **T** to 0, we have:

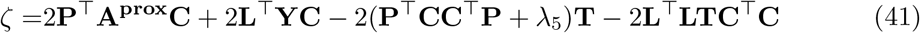

Using the Karush–Kuhn–Tucker conditions for the nonnegativity of **T**, we obtain:

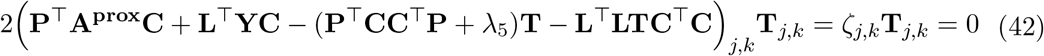

The parameter **T** is updated according:

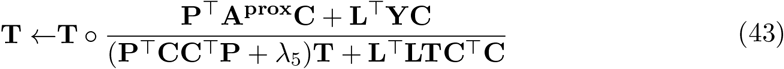

## 8. Update the community representation matrix R for EC features.

By fixing the parameters **C**, **T**, and **K**, we update **R**. The matrix **R** is updated such that the error is minimized:

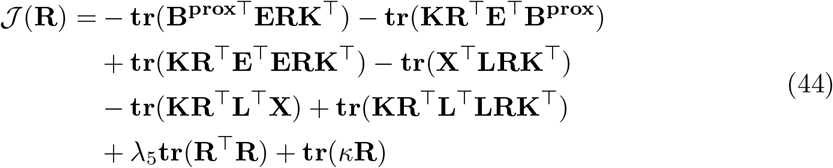

Taking the derivative of this error with respect to **R** to 0, we have:

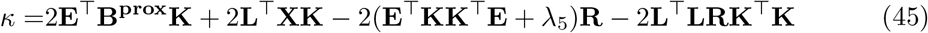

Using the Karush–Kuhn–Tucker conditions for the nonnegativity of **R**, we obtain:

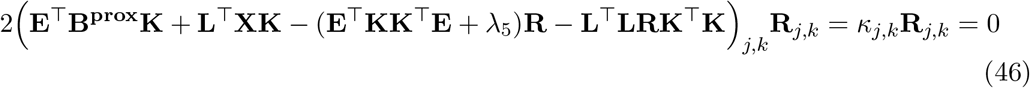

The parameter **R** is updated according:

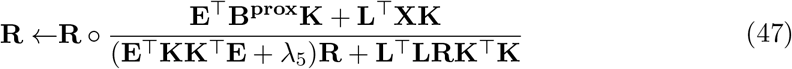

## 9. Update the weight matrix Θ.

By fixing the other parameters, we update Θ. The matrix Θ is updated such that the error is minimized:

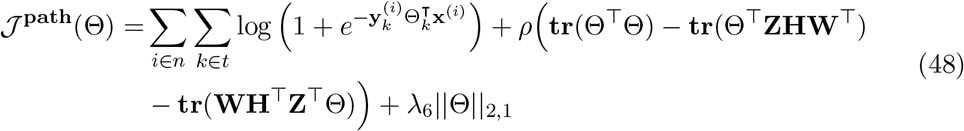

where *f* (.) is a non-lniear sigmoid function, i.e., 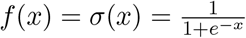. This choice can be generalized to any non-linear functions. By transforming **X** with *σ*(.) and Θ, our method enables pathway prediction. Taking the derivative of this error with respect to Θ to 0, we have:

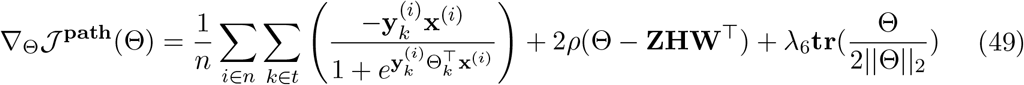

Due to non-closed form of the above equation, we use iterative gradient descent approach with a defined learning rate *η*. Hence, the general update rule for Θ becomes:

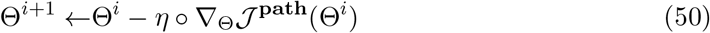

## 10. Update the auxiliary matrix L.

By fixing the rest of parameters in 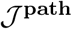, the matrix **L** is updated such that the error is minimized:

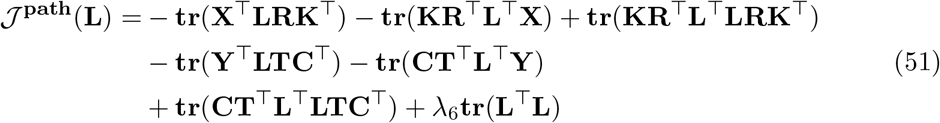

Taking the derivative of this error with respect to **L** to 0, we have:

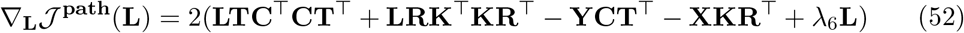

The parameter **L** is updated according:

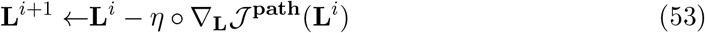

## 11. Update the auxiliary matrix Z.

By fixing the rest of parameters in 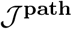, the matrix **Z** is updated such that the error is minimized:

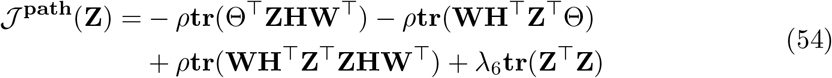

Taking the derivative of this error with respect to **Z** to 0, we have:

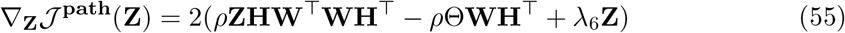

The parameter **Z** is updated according to gradient descent approach as:

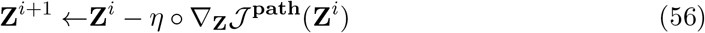

## 5.4 Appendix A4: Experimental Setup

In this section, we describe the experimental framework used to demonstrate triUMPF pathway prediction performance across multiple datasets spanning the genomic information hierarchy [25]. All experimental tests were conducted on a Linux server using 10 cores of Intel Xeon CPU E5-2650.

## 5.4.1 Association Matrices

MetaCyc v21 ([4]) was used to obtain the three association matrices, P2E (**M**), P2P, (**A**), and E2E (**B**). Some of the properties for each matrix are summarized in Table 3. All three matrices are extremely sparse. For example, **M** contains 2526 pathways, having an average of four EC associations per pathway, leaving more than 3600 columns with zero values. These matrices will be utilized to obtain higher-order proximity (Section 5.5.1) and to analyze triUMPF’s robustness (Section 5.5.2).

**Table 3:**
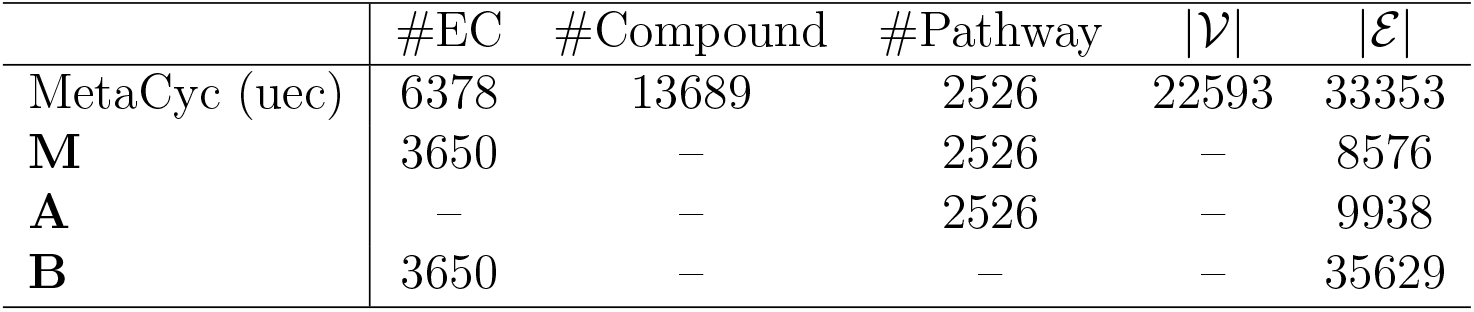
Characteristics of MetaCyc database and the three association matrices. MetaCyc (uec) denotes enzymatic reactions where links among enzymatic reactions are removed. The “–” indicates non applicable operation.

## 5.4.2 Description of Datasets

We report the performance of triUMPF using the following data: i)- T1 golden consisting of six PGDBs from the BioCyc collection (biocyc): *EcoCyc (v21)*, *HumanCyc (v19.5)*, *AraCyc (v18.5)*, *YeastCyc (v19.5)*, *LeishCyc (v19.5)*, and *TrypanoCyc (v18.5)*; ii)- three *E.coli* genomes consisting of E. coli K-12 substr. MG1655 (TAX-511145), E. coli str. CFT073 (TAX-199310), and E. coli O157:H7 str. EDL933 (TAX-155864) [36]; iii)- BioCyc (v20.5 T2 & 3) [5] consisting of 9255 Pathway/Genome Databases (PGDBs) with 1463 distinct pathways; iv)- reduced complexity of mealybug symbiont genomes from *Moranella* (GenBank NC-015735) and *Tremblaya* (GenBank NC-015736) encoding distributed metabolic pathways for amino acid biosynthesis [26]; v)- the Critical Assessment of Metagenome Interpretation (CAMI) initiative low complexity dataset (edwards.sdsu.edu/research/cami-challenge-datasets/), consisting of 40 genomes [31], and vi)- whole genome shotgun sequences from the Hawaii Ocean Time Series (HOTS) at 25m, 75m, 110m (sunlit) and 500m (dark) ocean depth intervals downloaded from the NCBI Sequence Read Archive under accession numbers SRX007372, SRX007369, SRX007370, SRX007371 [33]. T1 PGDBs were refined to include only those pathways that cross-intersect with the *MetaCyc* database (v21) [4]. The detailed characteristics of the datasets are summarized in Table 4. For each dataset 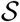, we use 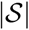 and 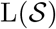 to represent the number of instances and pathway labels, respectively. In addition, we also present some characteristics of the multi-label datasets, which are denoted as:

1. Label cardinality 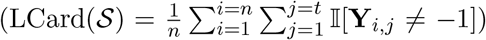 where 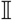 is an indicator function. It denotes the average number of pathways in 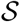.
2. Label density 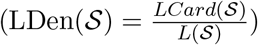. This is simply obtained through normalizing by the number 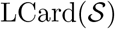 of total pathways in 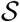.
3. Distinct labels 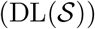. This notation indicates the number of distinct pathways in 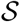.
4. Proportion of distinct labels 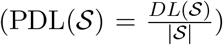. It represents the normalized version of 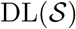, and is obtained by dividing DL(.) with the number of instances in 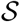.

The notations 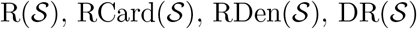, and 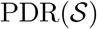 have similar meanings for the enzymatic reactions *ε* in 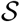. Finally, 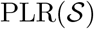 represents a ratio of 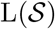 to 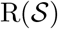.

## 5.4.3 Pathway and Enzymatic Reaction Features

triUMPF was trained using BioCyc v20.5 which contains less than 1460 trainable pathways. To offset this limit, we applied pathway2vec [24] using RUST-norm (or “crt”) module to obtain pathway and EC features, indicated by **P** and **E**, respectively, with the following settings: the number of memorized domain is 3, the explore and the in-out hyperparameters are 0.55 and 0.84, respectively, the number of sampled path instances was 100, the walk length is 100, the embedding dimension size was *m* = 128, the neighborhood size was 5, the size of negative samples was 5, and the used configuration of MetaCyc was “uec”, indicating links among ECs are being trimmed.

After generating node features, we only apply EC features to concatenate each example *i* according to:

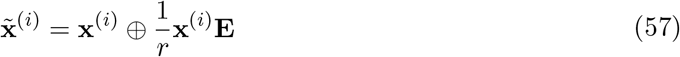

where ⊕ indicates the vector concatenation operation, **E** ∈ ℝ^*r*×*m*^ corresponds the feature matrix of ECs and *m* = 128. The addition of features results in a dimension of size *r* + *m*, where *r* = 3650. We expect by incorporating enzymatic reactions features into the original *r* dimensional example **x**^(*i*)^, the modified 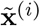 summarizes informative characteristics, which are expected to be useful in the prediction task.

**Table 4:**
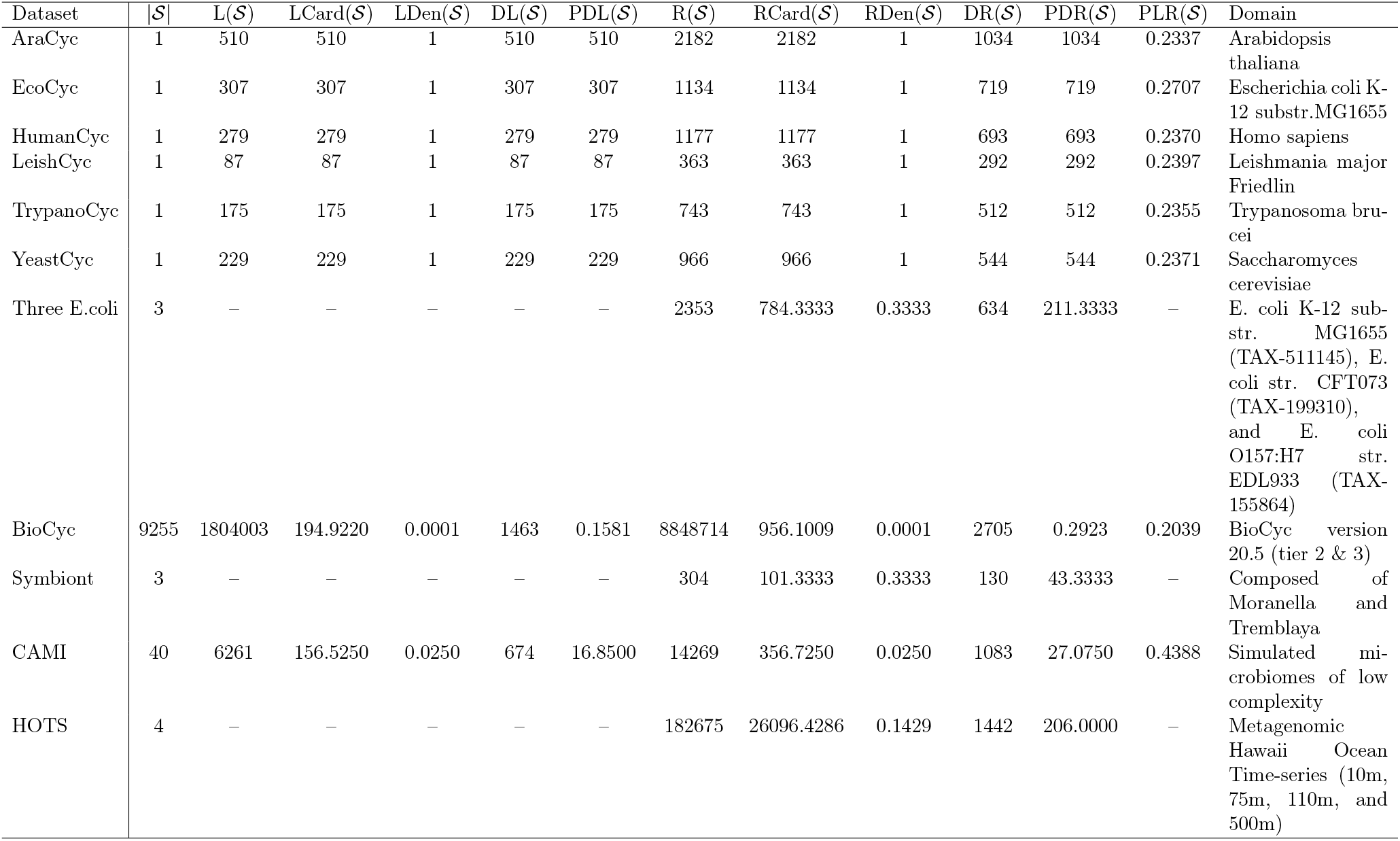
Experimental data set properties. The notations 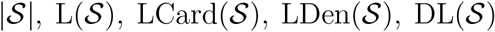 and 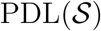 represent: number of instances, number of pathway labels, pathway labels cardinality, pathway labels density, distinct pathway labels, and proportion of distinct pathway labels for 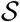, respectively. The notations 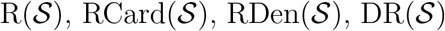 and 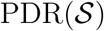 have similar meanings for the enzymatic reactions *ε* in 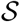. 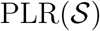 represents a ratio of 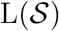 to 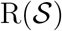 The last column denotes the domain of 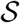.

## 5.4.4 Parameter Settings

For training, unless otherwise indicated, the learning rate was set to 0.0001, batch size to 50, number of epochs to 10, number of components *k* = 100, number of pathway and EC communities to *p* = 90 and *v* = 100, respectively. The higher-order proximity for **A^prox^** and **B^prox^**(corresponding P2P and E2E matrices, respectively, in Section 5.4.1) were set to *l^p^* = 3 and *l^e^* = 1 and their associated weights fixed as *ω* = 0.1 and *γ* = 0.3, respectively. The *α* and *β* were fixed to 10^9^. For the regularized hyperparameters *λ*_∗_, we performed 10-fold cross-validation on MetaCyc and a subsample of BioCyc T2 &3 data and found the settings *λ*_1:5_ = 0.01, *λ*_6_ = 10, and *ρ* = 0.001 to be optimum on golden T1 data.

## 5.5 Appendix A5: Experimental Results

Four tests were performed to benchmark the performance of triUMPF including parameter sensitivity, network reconstruction, impact of *ρ*, and metabolic pathway prediction.

## 5.5.1 Parameter Sensitivity

The impact of seven hyperparameters (*k, p, v, l_p_, l_e_*, *ω* and *γ*) was evaluated in relation to matrix reconstruction costs for (**M**, **A^prox^**, and **B^prox^**). The reconstruction cost (or error) defines the sum of mean squared errors accounted in the process of transforming the decomposed matrices into its original form where lower cost entails the decomposed low dimensional matrices were able to better capture the representations of the original matrix. We specifically evaluated the effects of varying the following parameters: i)- the number of components *k* ∈ {20, 50, 70, 90, 120}, ii)- the community size of pathway *p* ∈ {20, 50, 70, 90, 100} and EC *v* ∈ {20, 50, 70, 90, 100}, iii)- the higher-order proximity *l_p_* and *l_e_* ∈ {1, 2, 3}, and iv)- weights of the polynomial order *ω* and *γ* ∈ {0.1, 0.2, 0.3}. We used the full matrix **M**, for each test, however, for community detection, we used BioCyc T2 &3 data that is divided into training (80%), validation (5%) and test sets (15%). The final costs for community detection are reported based on the test set after 10 successive trials. In addition, we contrast triUMPF with the standard NMF for monitoring the reconstruction costs of **M** by varying *k* values. We emphasize that **M**, **A^prox^**, and **B^prox^** were collected from MetaCyc (Section 5.4.1) and not from BioCyc T2 &3 (Section 5.4.2).

Fig. 4 shows the effect of rank *k* on triUMPF performance. In general, we observe steady performance with increasing *k*. Although this contrasts standard NMF, where reconstruction cost decreases as the number of features increases it is expected because, unlike standard NMF, triUMPF exploits two types of correlations to recover **M**: i)- within ECs or pathways and ii)- betweenness interactions that serve as additional regularizers. As observed in Fig. 4, higher *k* values result in improved outcomes. Consequently, we selected *k* = 100 for downstream testing.

**Figure 4:**
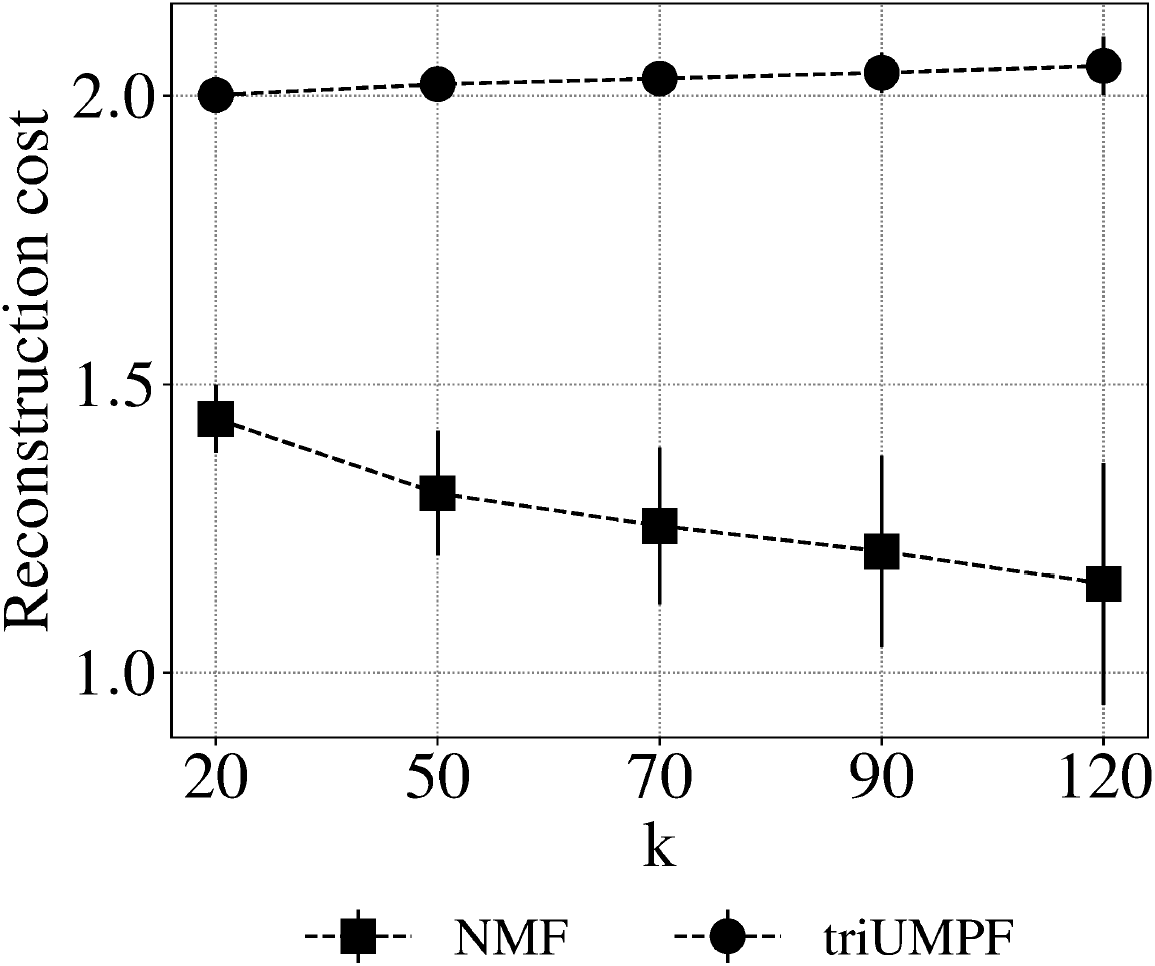
Sensitivity of components *k* based on reconstruction cost.

For community detection, we observed optimal results with respect to pathway community size at *p* = 20 under parameter settings *k* = 100 and *v* = 100, as shown in Fig. 5a. However, because **A^prox^** is so sparse, we suggest that this low rank may not correspond to the optimum community size. As with all methods of community detection triUMPF is sensitive to community size and requires empirical testing. Therefore, we tested settings between *p* = 20 and *p* = 100 and observed a decrease in performance under parameter settings *k* = 100 and *v* = 100 with *p* = 90 providing a balance between cost and increased community size. A similar result was observed for EC community size at *v* = 100 under parameter settings *p* = 90 and *k* = 100 in Fig. 5b.

**Figure 5:**
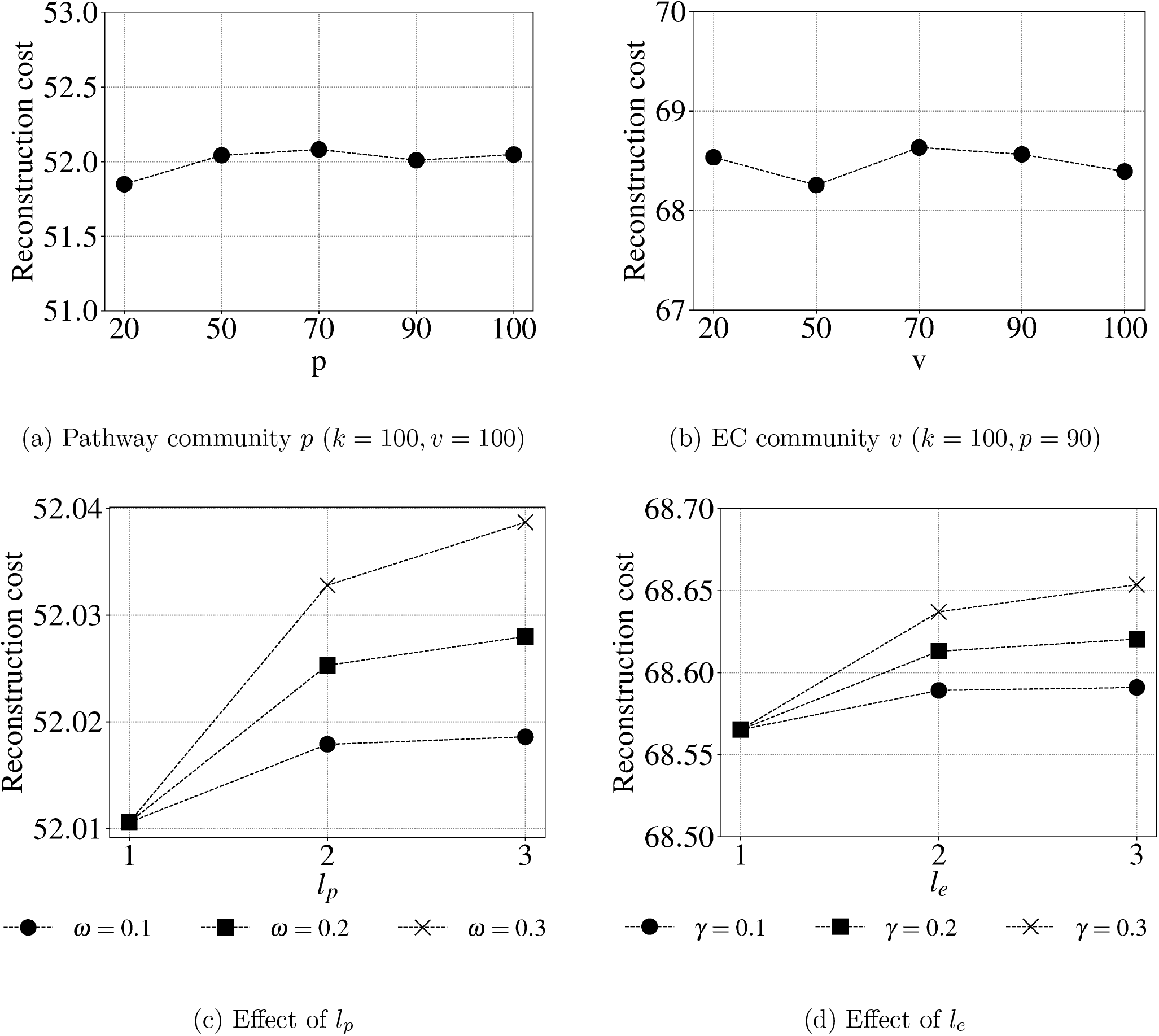
Sensitivity of community size and higher order proximity with weights based on reconstruction cost.

Finally, we show the effect of changing polynomial orders, and their weights on triUMPF performance. From Fig. 5c, we see that reconstruction cost progressively increases with varying higher orders for *l_p_* for all the three weights *ω*. However, for the same reasons described above, we prefer more long distances with less weight to preserve community structure, and remarkably, when *ω* = 0.1 triUMPF performance was relatively stable after the second order. The same conclusion can be drawn for *l_e_* and its associated weights *γ* in Fig. 5d.

Based on these results, triUMPF performance is stable while minimizing cost under the following parameter settings: *k* = 100, *p* > 90, *e* > 90, *l_p_* = 3, *ω* = 0.1, *l_e_* = 1, and *γ* = 0.3. Therefor, we recommend these settings for both MetaCyc and BioCyc T2 &3.

## 5.5.2 Network Reconstruction

In this section, we explore the robustness of triUMPF when exposed to noise. Links were randomly removed from **M**, **A**, and **B** according to *ε* ∈ {20%, 40%, 60%, 80%}. We used the partially linked matrices to refine parameters while comparing the reconstruction cost against the full association matrices **M**, **A** and **B**. Specifically for **M**, we varied components of **M** according to *k* ∈ {20, 50, 70, 90, 120} along with *ϵ*. For all experiments, both MetaCyc and BioCyc T2 &3 were applied for training using hyperparameters described in Section 3.4 of the primary text.

Fig. 6a indicate that by progressively increasing noise *ε* to **M**, the reconstruction cost increases when *k* is low. As more features are incorporated the cost at all noise levels steadily decreases up to *k* = 100. This tendency indicates that both pathway and EC features (**P** and **E** contain useful correlations that contribute to the resilience of triUMPF’s performance when **M** is perturbed.

**Figure 6:**
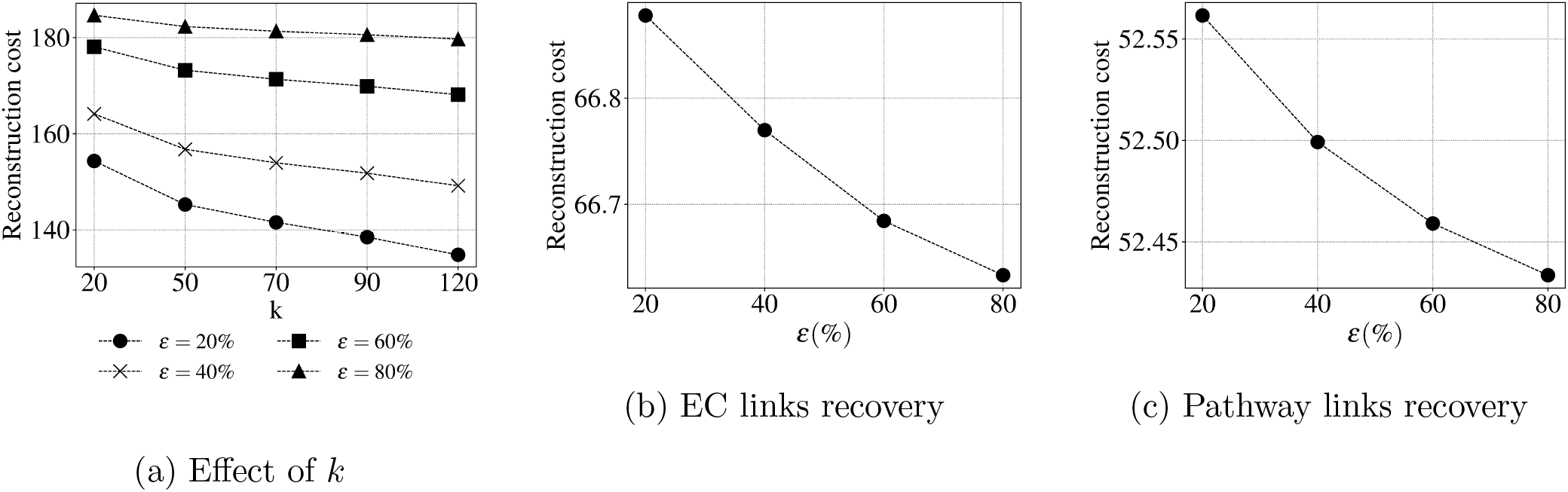
Link prediction results by varying noise levels *ε* ∈ {20%, 40%, 60%, 80%} based on reconstruction cost.

For **A^prox^** and **B^prox^**, as shown in Figs 6b and 6b, the costs are reduced in the presence of noise, which is not surprising as the reconstruction of associated communities are constrained on both data and **A^prox^** and **B^prox^**. These results are directly linked to the sparseness of both matrices, as previously described in [8]. The pathway graph network, depicted in Fig. 1 of the primary text, indicates that many pathways constitute islands with no direct links, while some pathways are densely connected. For community detection, it is sufficient to group nodes that are densely connected, while links between communities can remain sparse. The same line of reasoning follows for the EC network.

## 5.5.3 Impact of *ρ*

Fig. 7 shows the inverse effect in predictive performance on T1 golden datasets when decreasing *ρ* before reaching a performance plateau at *ρ* = 0.001. The hyperparameter *ρ* in Eq. 5 controls the amount of information propagation from **M** to pathway label coefficients Θ. This suggests, in practice, lesser constraints should be emphasized on Θ, while not neglecting associations between EC numbers and pathways indicated in **M**.

**Figure 7:**
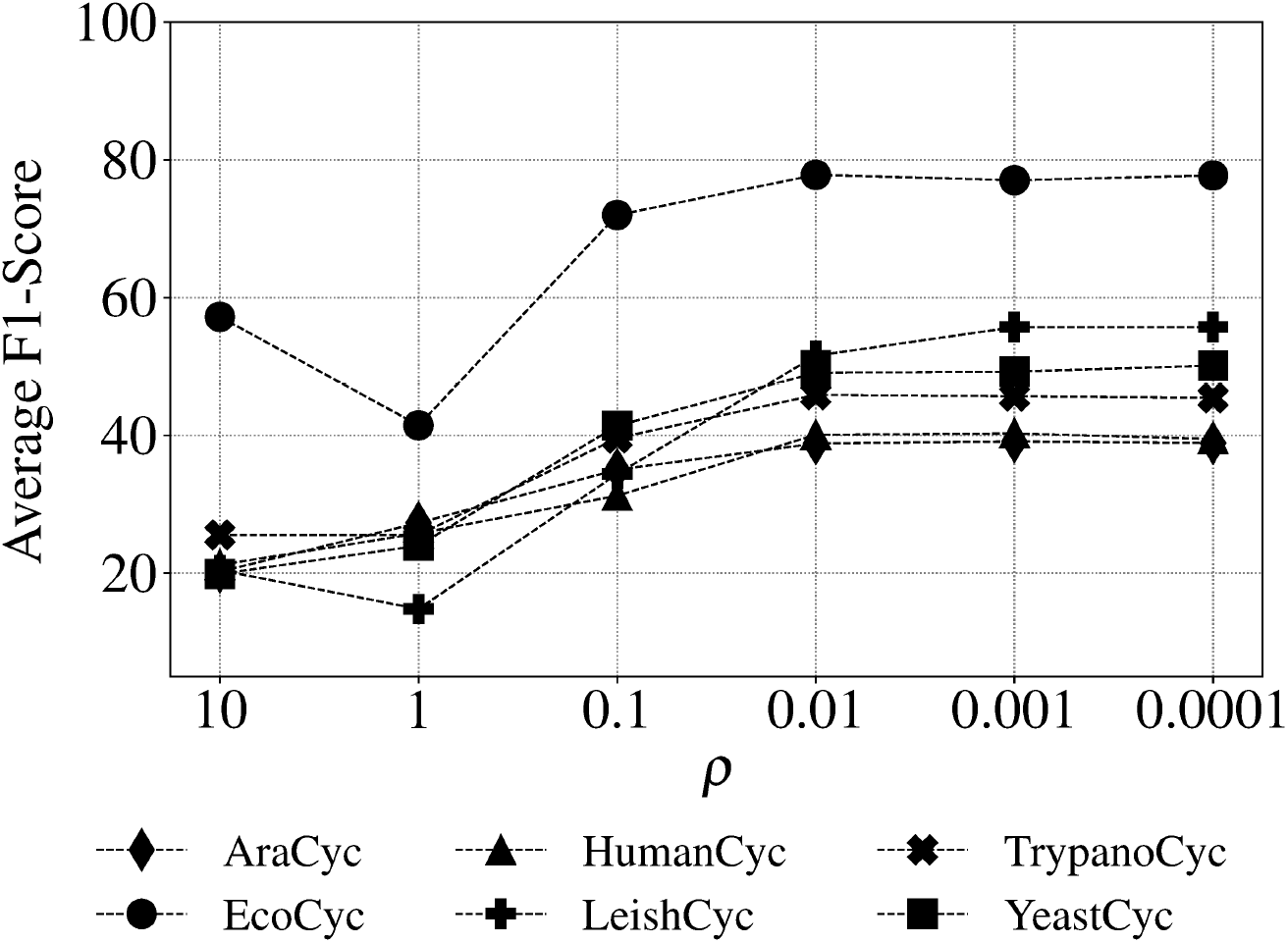
Effect of *ρ* based on average F1 score using golden datasets.

## 5.5.4 Metabolic Pathway Prediction

Here, we investigate the effectiveness of triUMPF for the pathway prediction task on i)- T1 golden data, ii)- three *E. coli* data, and iii)- HOTS.

## T1 Golden Data.

We compare the performance of triUMPF on 6 benchmark datasets, as described in Section 5.4.2, against the other pathway prediction algorithms using four evaluation metrics: *Hamming loss*, *average precision*, *average recall*, and *average F1 score*. As shown in Table 5, triUMPF achieved competitive performance against the other methods in terms of average precision.

**Table 5:** Predictive performance of each comparing algorithm on 6 golden T1 data. For each performance metric, ‘↓’ indicates the smaller score is better while ‘↑’ indicates the higher score is better.

| Methods | Hamming Loss ↓ |  |  |  |  |  |
| --- | --- | --- | --- | --- | --- | --- |
|  | EcoCyc | HumanCyc | AraCyc | YeastCyc | LeishCyc | TrypanoCyc |
| PathoLogic | 0.0610 | **0.0633** | 0.1188 | **0.0424** | **0.0368** | **0.0424** |
| MinPath | 0.2257 | 0.2530 | 0.3266 | 0.2482 | 0.1615 | 0.2561 |
| mlLGPR | 0.0804 | **0.0633** | **0.1069** | 0.0550 | 0.0380 | 0.0590 |
| triUMPF | **0.0435** | 0.0954 | 0.1560 | 0.0649 | 0.0443 | 0.0776 |
| Methods | Average Precision ↑ |  |  |  |  |  |
|  | EcoCyc | HumanCyc | AraCyc | YeastCyc | LeishCyc | TrypanoCyc |
| PathoLogic | 0.7230 | **0.6695** | 0.7011 | 0.7194 | **0.4803** | **0.5480** |
| MinPath | 0.3490 | 0.3004 | 0.3806 | 0.2675 | 0.1758 | 0.2129 |
| mlLGPR | 0.6187 | 0.6686 | 0.7372 | 0.6480 | 0.4731 | 0.5455 |
| triUMPF | **0.8662** | 0.6080 | **0.7377** | **0.7273** | 0.4161 | 0.4561 |
| Methods | Average Recall ↑ |  |  |  |  |  |
|  | EcoCyc | HumanCyc | AraCyc | YeastCyc | LeishCyc | TrypanoCyc |
| PathoLogic | 0.8078 | 0.8423 | 0.7176 | 0.8734 | 0.8391 | 0.7829 |
| MinPath | **0.9902** | **0.9713** | **0.9843** | **1.0000** | **1.0000** | **1.0000** |
| mlLGPR | 0.8827 | 0.8459 | 0.7314 | 0.8603 | 0.9080 | 0.8914 |
| triUMPF | 0.7590 | 0.3835 | 0.3529 | 0.3319 | 0.7126 | 0.6229 |
| Methods | Average F1 ↑ |  |  |  |  |  |
|  | EcoCyc | HumanCyc | AraCyc | YeastCyc | LeishCyc | TrypanoCyc |
| PathoLogic | 0.7631 | 0.7460 | 0.7093 | **0.7890** | 0.6109 | 0.6447 |
| MinPath | 0.5161 | 0.4589 | 0.5489 | 0.4221 | 0.2990 | 0.3511 |
| mlLGPR | 0.7275 | **0.7468** | **0.7343** | 0.7392 | **0.6220** | **0.6768** |
| triUMPF | **0.8090** | 0.4703 | 0.4775 | 0.4735 | 0.5254 | 0.5266 |

**Table 6:**
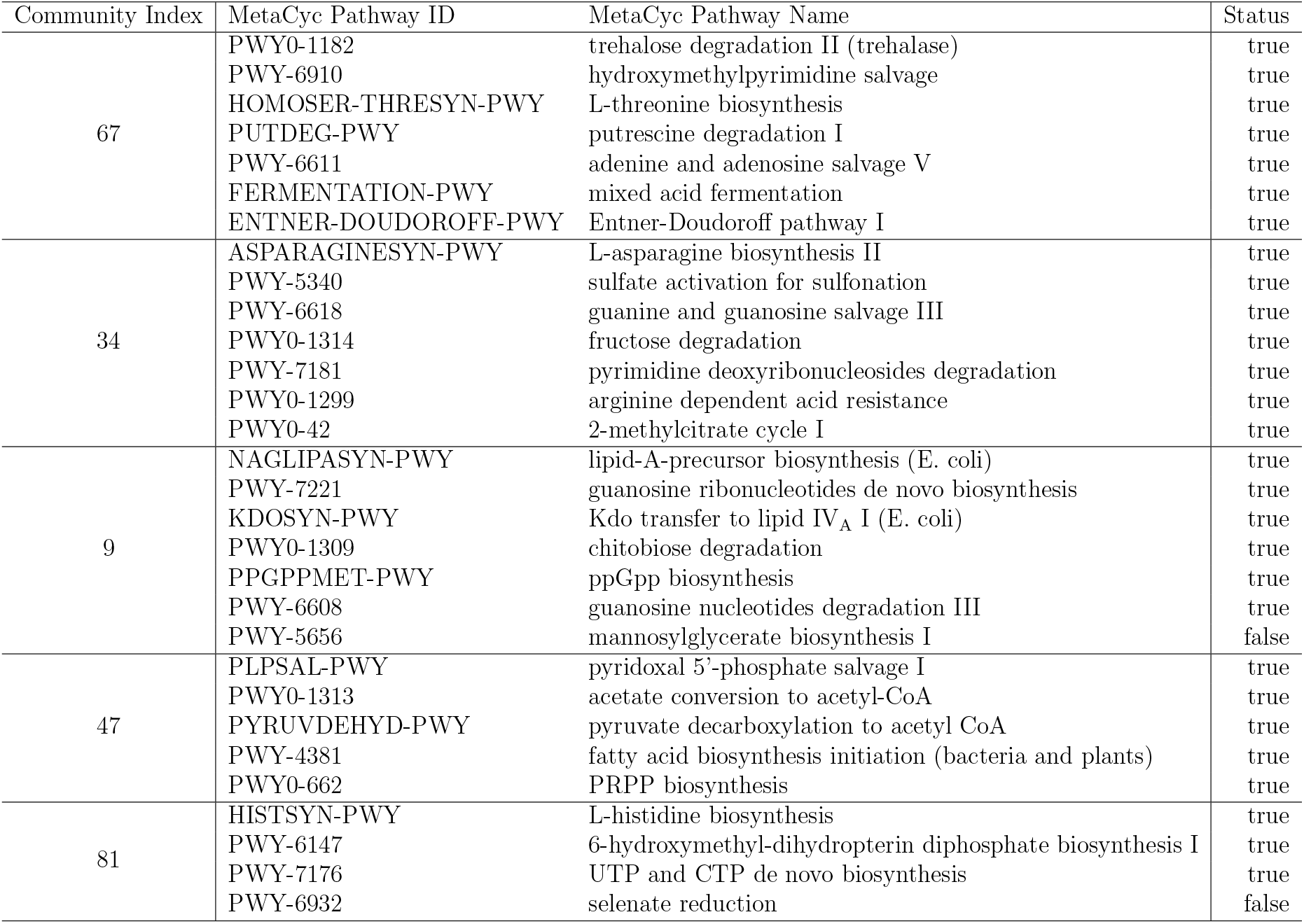
Top 5 communities with pathways predicted by triUMPF for E. coli K-12 substr. MG1655 (TAX-511145). The last column asserts whether a pathway is present in or absent (a false-positive pathway) from EcoCyc reference data.

| Community Index | MetaCyc Pathway ID | MetaCyc Pathway Name | Status |
| --- | --- | --- | --- |
| 67 | PWY0-1182 | trehalose degradation II (trehalase) | true |
|  | PWY-6910 | hydroxymethylpyrimidine salvage | true |
|  | HOMOSER-THRESYN-PWY | L-threonine biosynthesis | true |
|  | PUTDEG-PWY | putrescine degradation I | true |
|  | PWY-6611 | adenine and adenosine salvage V | true |
|  | FERMENTATION-PWY | mixed acid fermentation | true |
|  | ENTNER-DOUDOROFF-PWY | Entner-Doudoroff pathway I | true |
| 34 | ASPARAGINESYN-PWY | L-asparagine biosynthesis II | true |
|  | PWY-5340 | sulfate activation for sulfonation | true |
|  | PWY-6618 | guanine and guanosine salvage III | true |
|  | PWY0-1314 | fructose degradation | true |
|  | PWY-7181 | pyrimidine deoxyribonucleosides degradation | true |
|  | PWY0-1299 | arginine dependent acid resistance | true |
|  | PWY0-42 | 2-methylcitrate cycle I | true |
| 9 | NAGLIPASYN-PWY | lipid-A-precursor biosynthesis (E. coli) | true |
|  | PWY-7221 | guanosine ribonucleotides de novo biosynthesis | true |
|  | KDOSYN-PWY | Kdo transfer to lipid IV<sub>A</sub> I (E. coli) | true |
|  | PWY0-1309 | chitobiose degradation | true |
|  | PPGPPMET-PWY | ppGpp biosynthesis | true |
|  | PWY-6608 | guanosine nucleotides degradation III | true |
|  | PWY-5656 | mannosylglycerate biosynthesis I | false |
| 47 | PLPSAL-PWY | pyridoxal 5'-phosphate salvage I | true |
|  | PWY0-1313 | acetate conversion to acetyl-CoA | true |
|  | PYRUVDEHYD-PWY | pyruvate decarboxylation to acetyl CoA | true |
|  | PWY-4381 | fatty acid biosynthesis initiation (bacteria and plants) | true |
|  | PWY0-662 | PRPP biosynthesis | true |
| 81 | HISTSYN-PWY | L-histidine biosynthesis | true |
|  | PWY-6147 | 6-hydroxymethyl-dihydropterin diphosphate biosynthesis I | true |
|  | PWY-7176 | UTP and CTP de novo biosynthesis | true |
|  | PWY-6932 | selenate reduction | false |

**Table 7:** 18 amino acid biosynthesis pathways and 27 pathway variants.

| Amino Acid | MetaCyc Pathway ID | MetaCyc Pathway Name |
| --- | --- | --- |
| Arginine | ARGSYNBSUB-PWY | L-arginine biosynthesis II (acetyl cycle) |
|  | PWY-5154 | L-arginine biosynthesis III (via N-acetyl-L-citrulline) |
|  | PWY-7400 | L-arginine biosynthesis IV (archaeobacteria) |
| Asparagine | ASPARAGINE-BIOSYNTHESIS | L-asparagine biosynthesis I |
|  | ASPARAGINESYN-PWY | L-asparagine biosynthesis II |
| Chorismate | PWY-6163 | chorismate biosynthesis from 3-dehydroquinate |
| Cysteine | CYSTSYN-PWY | L-cysteine biosynthesis I |
|  | PWY-6308 | L-cysteine biosynthesis II (tRNA-dependent) |
| Glutamine | GLNSYN-PWY | L-glutamine biosynthesis I |
| Glycine | GLYSYN-PWY | glycine biosynthesis I |
|  | GLYSYN-THR-PWY | glycine biosynthesis IV |
| Histidine | HISTSYN-PWY | L-histidine biosynthesis |
| Isoleucine | ILEUSYN-PWY | L-isoleucine biosynthesis I (from threonine) |
|  | PWY-5104 | L-isoleucine biosynthesis IV |
| Leucine | LEUSYN-PWY | L-leucine biosynthesis |
| Lysine | DAPLYSINESYN-PWY | L-lysine biosynthesis I |
|  | PWY-2941 | L-lysine biosynthesis II |
|  | PWY-2942 | L-lysine biosynthesis III |
| Methionine | HOMOSER-METSYN-PWY | L-methionine biosynthesis I |
|  | PWY-702 | L-methionine biosynthesis II |
| Phenylalanine | PHESYN | L-phenylalanine biosynthesis I |
| Proline | PROSYN-PWY | L-proline biosynthesis I |
| Serine | SERSYN-PWY | L-serine biosynthesis |
| Threonine | HOMOSER-THRESYN-PWY | L-threonine biosynthesis |
| Tryptophan | TRPSYN-PWY | L-tryptophan biosynthesis |
| Tyrosine | TYRSYN | L-tyrosine biosynthesis I |
| Valine | VALSYN-PWY | L-valine biosynthesis |

## Three E.coli Data.

Fig. 8 shows pathway communities observed for MG1655, CFT073 and EDL933 using BioCyc T2 &3 including MetaCyc in training. Fig. 9 shows that PathoLogic was able to infer over 90 additional pathways when taxonomic pruning is disabled. Table 7 summarizes GapMind [29] results for MG1655, CFT073 and EDL933. Fig. 10 shows the results for both PathoLogic with taxonomic pruning enabled and triUMPF. Without taxonomic pruning, PathoLogic predicted 56 pathways across the three strains encompassing 15 amino acid biosynthesis pathways and 20 pathway variants, including *L-proline biosynthesis II (from arginine)* pathway that is known only for eukaryotes (Fig. 11), consequently, increasing false-positive pathway prediction.

**Figure 8:**
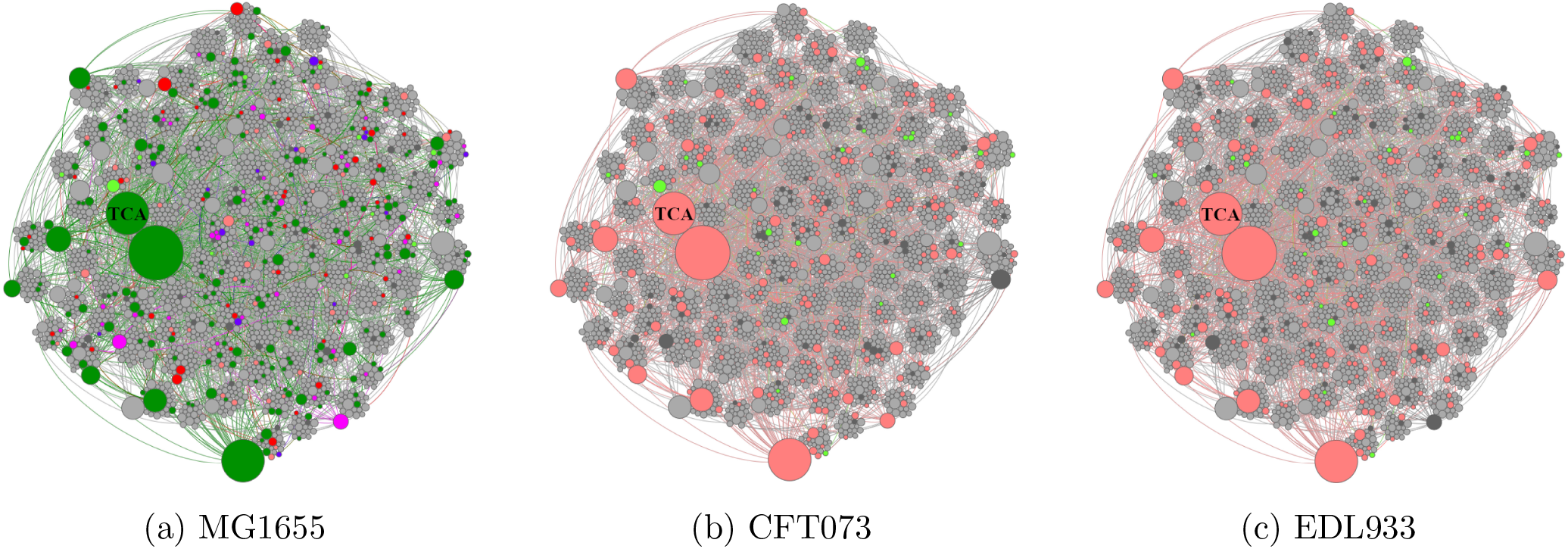
Pathway community networks for related T1 and T3 organismal genomes. Pathway communities for (a) E. coli K-12 substr. MG1655 (TAX-511145), (b) E. coli str. CFT073 (TAX-199310), and (c) E. coli O157:H7 str. EDL933 (TAX-155864) based on community detection. Nodes colored in *dark grey* indicate pathways predicted by PathoLogic; *lime* pathways predicted by triUMPF; *salmon* pathways predicted by both PathoLogic and triUMPF; *red* expected pathways not predicted by both PathoLogic and triUMPF; *magenta* expected pathways predicted only by PathoLogic; *purple* expected pathways predicted solely by triUMPF; and *green* expected pathways predicted by both PathoLogic and triUMPF. *light-grey* indicates pathways not expected to be encoded in either organismal genome. The node sizes reflect the degree of associations between pathways.

**Figure 9:**
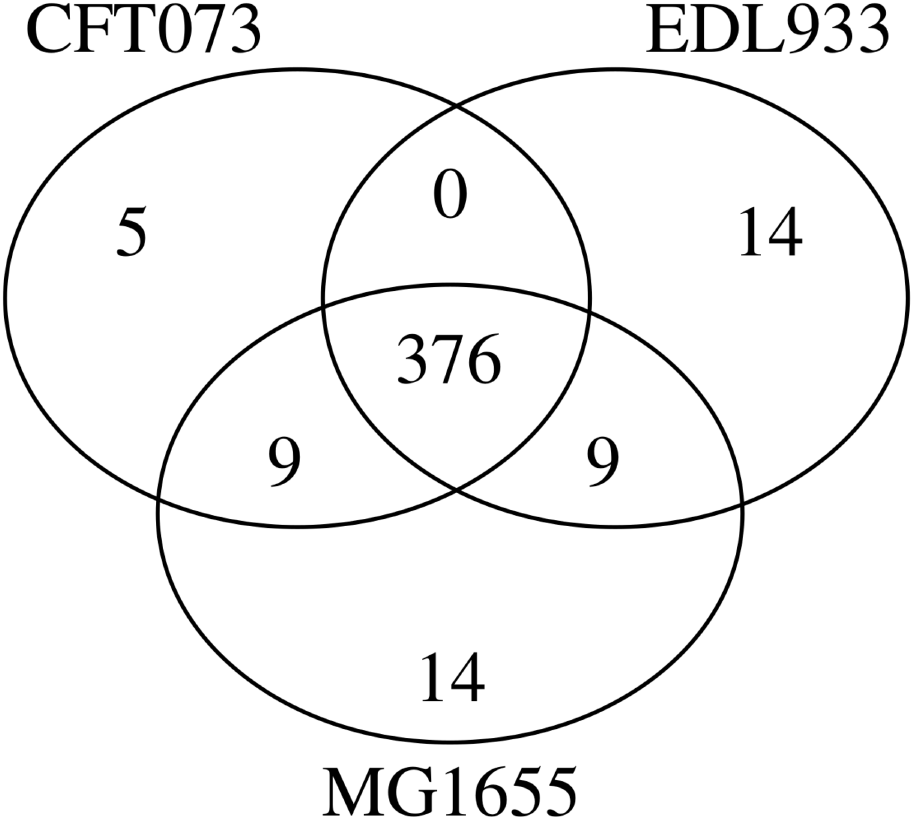
A three way set analysis of predicted pathways for E. coli K-12 substr. MG1655 (TAX-511145), E. coli str. CFT073 (TAX-199310), and E. coli O157:H7 str. EDL933 (TAX-155864) using PathoLogic (without taxonomic pruning).

**Figure 10:**
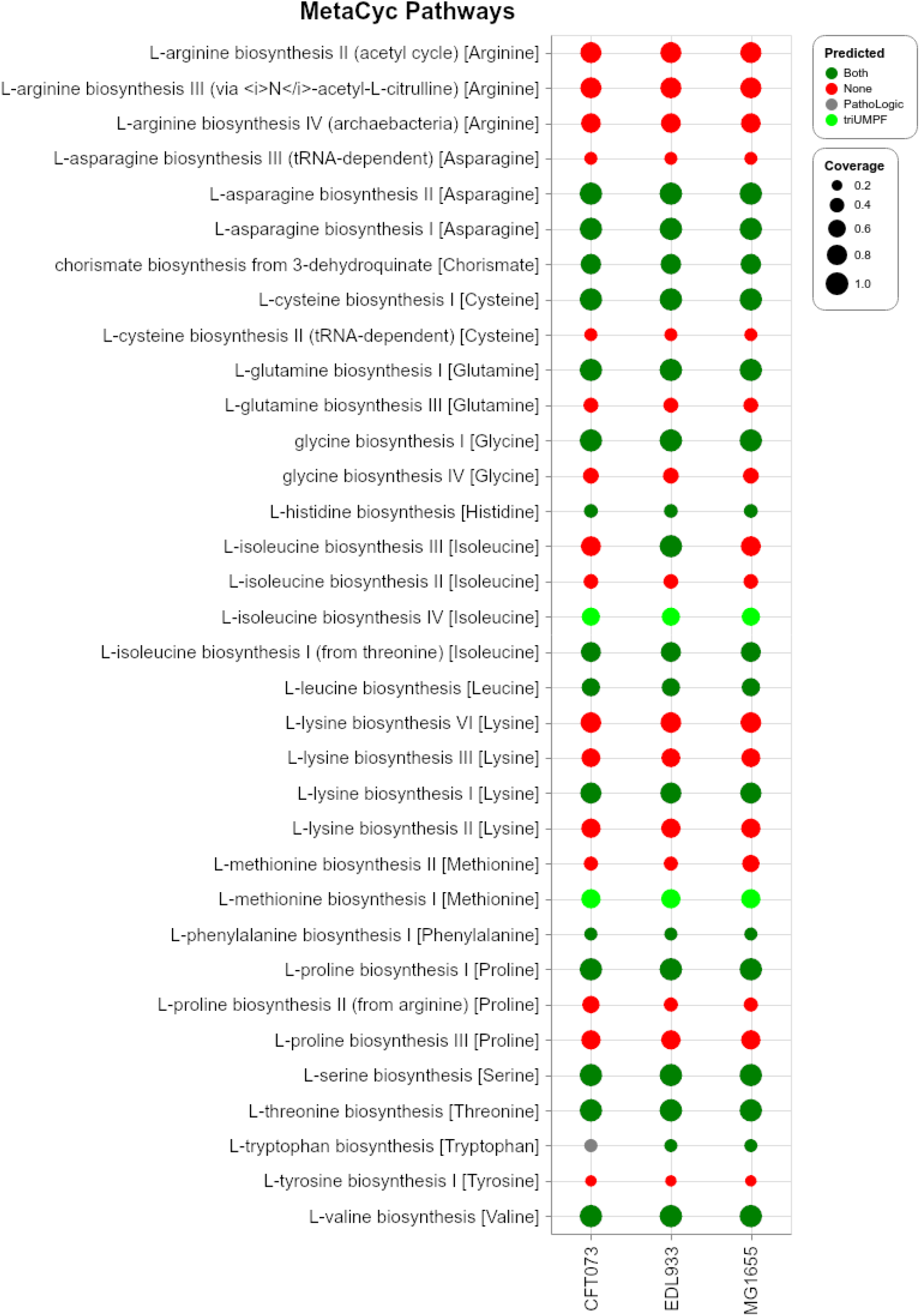
Comparison of predicted pathways for E. coli K-12 substr. MG1655 (TAX-511145), E. coli str. CFT073 (TAX-199310), and E. coli O157:H7 str. EDL933 (TAX-155864) datasets between PathoLogic (taxonomic pruning) and triUMPF. Red circles indicate that neither method predicted a specific pathway while green circles indicate that both methods predicted a specific pathway. Lime circles indicate pathways predicted solely by mlLGPR and gray circles indicate pathways solely predicted by PathoLogic.The size of circles corresponds to associated pathway coverage information.

**Figure 11:**
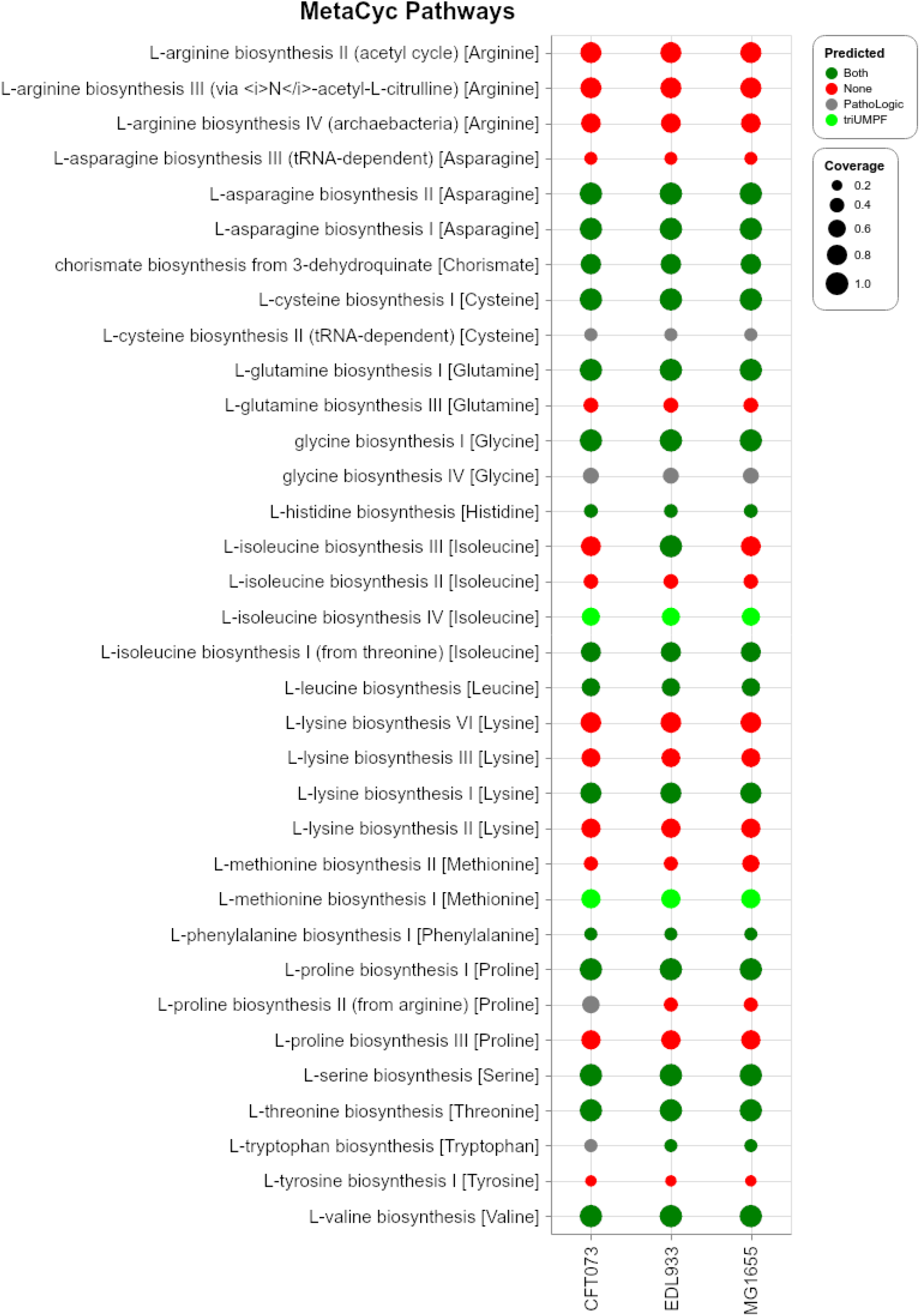
Comparison of predicted pathways for E. coli K-12 substr. MG1655 (TAX-511145), E. coli str. CFT073 (TAX-199310), and E. coli O157:H7 str. EDL933 (TAX-155864) datasets between PathoLogic (without taxonomic pruning) and triUMPF. Red circles indicate that neither method predicted a specific pathway while green circles indicate that both methods predicted a specific pathway. Lime circles indicate pathways predicted solely by mlLGPR and gray circles indicate pathways solely predicted by PathoLogic. The size of circles corresponds the associated coverage information.

## HOTS water column.

Here, we use triUMPF to infer a set of pathways from the HOTS water column spanning sunlit and dark ocean depth intervals comparing results to other prediction methods including PathoLogic and mlLGPR. The results are presented in Fig. 12.

**Figure 12:**
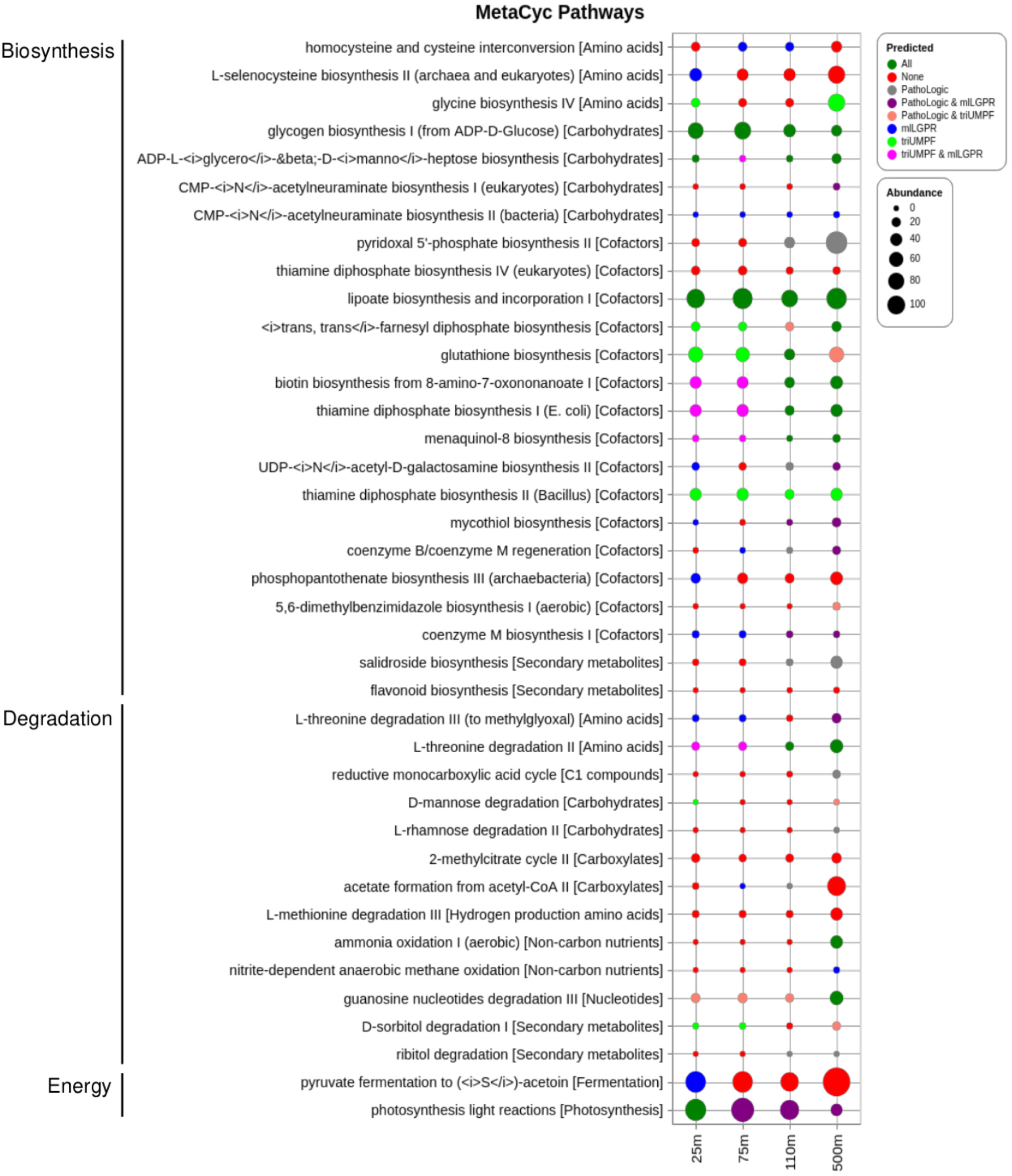
Comparative study of predicted pathways for HOT DNA samples. The size of circles corresponds the associated coverage information.

## Availability of Data and Materials

The triUMPF source code is available under the MIT License on GitHub (hallamlab/triUMPF) with detailed descriptions on how to install and execute all commands run to generate results in our GitHub repository. The MetaCyc database can be obtained from metacyc.org. The T1 golden datasets can be downloaded from biocyc.org. For the symbiotic *Candidatus Moranella endobia* and *Candidatus Tremblaya princeps* genomes, they can be downloaded from GenBank under accession numbers NC-015735 and NC-015736 while the simulated CAMI low complexity dataset can be obtained from edwards.sdsu.edu/research/cami-challenge-datasets. Unassembled whole genome shotgun DNA pyrosequences from HOTS (10m, 75m, 110m, and 500m) can be obtained from the NCBI Sequence Read Archive under accession numbers SRX007372, SRX007369, SRX007370, SRX007371. The preprocessed datasets used in this paper can be downloaded from zenodo.org/YLIBG lKhPZ. The same zenodo repo contains a pre-trained tri-UMPF (triUMPF Xe.pkl) using configurations stated in Section 5.4. We also included the three preprocessed E.coli data in the github repo under the “sample” directory.

## Acknowledgments

We would like to thank Connor Morgan-Lang, Julia Anstett, Kishori Konwar and Aria Hahn for lucid discussions on the function of the triUMPF model and all members of the Hallam Lab for helpful comments along the way.

